# Bayesian Linear Mixed Models for Motif Activity Analysis

**DOI:** 10.1101/782615

**Authors:** Simone Lederer, Tom Heskes, Simon J. van Heeringen, Cornelis A. Albers

## Abstract

**Motivation:** Cellular identity and behavior is controlled by complex gene regulatory networks. Transcription factors (TFs) bind to specific DNA sequences to regulate the transcription of their target genes. On the basis of these TF motifs in cis-regulatory elements we can model the influence of TFs on gene expression. In such models of TF motif activity the data is usually modeled assuming a linear relationship between the motif activity and the gene expression level. A commonly used method to model motif influence is based on Ridge Regression. One important assumption of linear regression is the independence between samples. However, if samples are generated from the same cell line, tissue, or other biological source, this assumption may be invalid. This same assumption of independence is also applied to different, yet similar, experimental conditions, which may also be inappropriate. In theory, the independence assumption between samples could lead to loss in signal detection. Here we investigate whether a Bayesian model that allows for correlations results in more accurate inference of motif activities.

**Results:** We extend the Ridge Regression to a Bayesian Linear Mixed Model, which allows us to model dependence between different samples. In a simulation study, we investigate the differences between the two model assumptions. We show that our Bayesian Linear Mixed Model implementation outperforms Ridge Regression in a simulation scenario where the noise, the signal that can not be explained by TF motifs, is uncorrelated. However, we demonstrate that there is no such gain in performance if the noise has a similar covariance structure over samples as the signal that can be explained by motifs. We give a mathematical explanation to why this is the case. Using two representative real data sets we show that at most *∼* 40% of the signal is explained by motifs using the linear model. With these data there is no advantage to using the Bayesian Linear Mixed Model, due to the similarity of the covariance structure.

**Availability & Implementation:** The project implementation is available at https://github.com/Sim19/SimGEXPwMotifs.

## Introduction

Cell type-specific gene expression programs are mainly driven by differential expression and binding of transcription factors (TFs). The human genome contains *∼* 1, 600 TFs, which represent 8% of all genes (Lambert *et al*., 2018). These proteins bind DNA in a sequence-specific manner and typically have a 1000-fold or greater preference for their cognate binding site as compared to other sequences (Geertz *et al*., 2012). By binding to cis-regulatory regions, promoters and enhancers, they can control the chromatin environment and the expression of downstream target genes (Lambert *et al*., 2018). Cell type identity is determined by the expression of a select number of TFs. This is evidenced by the growing number of cell reprogramming protocols that rely on the activation of a few TFs to reprogram the cell state, for instance from a somatic cell to a pluripotent stem cell (Lee and Young, 2013; Takahashi and Yamanaka, 2006). Mis-regulation of TF expression or binding is associated with a variety of diseases, such as developmental disorders and cancer (Lee and Young, 2013). Hence, it is of great importance to understand the mechanisms of gene regulation driven by TFs.

TFs bind to specific DNA sequences called sequence motifs. These motifs are relatively short, with a length usually ranging from six to twelve nucleotides, and flexible in the sense that several TFs can bind to the same motif (Lambert *et al*., 2018). The binding sites of TFs can be determined genome-wide using chromatin immunoprecipitation with specific antibodies followed by high-throughput sequencing (ChIP-seq). Although ChIP-seq studies suggest that many TF binding events appear to be not functional, the presence of a sequence motif is still predictive of gene expression (Lambert *et al*., 2018). With a linear regression model, in which the sequence information is used to model gene expression, one can learn the TFs that play a major role in gene regulation (The Fantom Consortium and the Riken Omics Science Center, 2009; Balwierz *et al*., 2014; Osmanbeyoglu *et al*., 2014; Schmidt *et al*., 2017; Madsen *et al*., 2018). Typical approaches either use linear regression with *ℒ*^2^-regularization (Ridge Regression) or a combination of *ℒ*^1^ and *ℒ*^2^ -regularization (ElasticNet). These approaches tend to explain only a small fraction of the variation of gene expression. However, due to the large number of genes, the coefficients are generally highly significant and can be interpreted as a measure of transcription factor activity.

One of the key assumptions of a linear regression model is the independence between samples. If samples originate from the same cell line, tissue or other biological source, this assumption may be invalid. In addition, related cell types will also have similar gene expression profiles, where the expression of many genes will be highly correlated.

Here, we propose a Bayesian Linear Mixed Model that builds upon the previously described Bayesian Ridge Regression (The Fantom Consortium and the Riken Omics Science Center, 2009; Balwierz *et al*., 2014), but allows for correlated motif activity between samples. Our model relaxes the rigid independence assumption common to earlier approaches.

We compare our full Bayesian Linear Mixed Model with Bayesian Ridge Regression on simulated data, for which we control the degree of correlation between samples. We show that the Bayesian Linear Mixed Model formulation outperforms the Ridge Regression for data with randomly distributed noise. This is the case especially for highly correlated data. We further show that the Bayesian Linear Mixed Model loses its superiority over the Ridge Regression if only a small part of the gene expression signal can be explained by motif influence, while other influential factors contribute largely to the gene expression. We confirm the observations made during the simulation study on two real-world datasets, in which a significant amount of the biological signal cannot be uniquely explained by a linear relationship of motifs. We can explain this phenomenon mathematically and give more technical details regarding the computation of the model.

## Methods

In this section we introduce the mathematical models that are used throughout this paper. First, we introduce the linear model that represents the signal of expression data **Y***_G,C_* as a linear combination of motif scores and their influential weights. Second, we present a Bayesian perspective on the model. And third, we show that the Ridge Regression formulation is a special case of the newly introduced Bayesian formulation. We then outline in detail how we simulate data, which we use to compare the complete Bayesian perspective and the Ridge Regression model.

In the following sections we will interchangeably use sample or condition.

**Inference of motif activities**

The general model used throughout this paper models gene expression **y**_*g,c*_ in condition *c* as a linear function of motif scores **m**_*t,g*_ of motifs *t* = {1, …, *T* } at the promoter region of gene *g*:

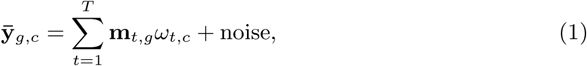

with the normalized gene expression data 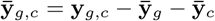, where 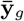 is the average signal over the promoter of gene *g* over all conditions *C* and 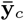 is the average signal of condition *c* over all genes *G*. In the following, we will refer to the normalized gene expression simply as **Y***_G,C_*. The term “noise” represents all signal that cannot be explained by the model, i.e. the linear combination of the motif scores **M***_T,G_*. This can be any technical noise, motif influence for which the linear assumption might be too simplistic, but also any other source that drives the gene expression **y**_*g,c*_ and is not modeled. The model was originally introduced by The Fantom Consortium and the Riken Omics Science Center (2009) and subsequently expanded by Balwierz *et al.* (2014).

For the application of the above model to the H3K27ac dataset, we re-use the above notation, where *G* represents the dimensionality in enhancer space (not gene space). Instead of computing motif scores **M***_T,G_* in the promoter region, we compute them in the enhancer region and the expression signal **Y***_G,C_* is the H3K27ac ChIP-seq signal.

### Bayesian Linear Mixed Models

The main idea behind a Bayesian formulation is to include prior knowledge about the data, called the prior. Here, we model *ω*_*t,c*_, the influence of motif *t* ∈ {1, …, *T* } in condition *c* ∈ {1, …, *C*}, as a normally distributed prior with mean zero, with *σ*^*2*^ **I***_T_* being the covariance over all motifs and **V***_C_* the covariance over all conditions:

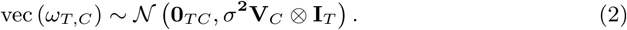

We use the vector notation for the matrix normal distribution, for which we make use of the Kronecker product between the covariance matrices. Another way of mathematically writing the model is *ℳ𝒩_T,C_*(**0***_T,C_, σ*^**2**^**I***_T_*, **V***_C_*). Note that in the vectorized notation the mean **0***_T,C_* is written in vector notation, too: vec (**0***_T,C_*) = **0***_TC_*.

In this paper, we assume independence between motifs. Extending the assumption to dependence between motifs with covariance matrix **Ψ** could easily be implemented. Combining the prior knowledge about *ω*_*T,C*_ in Eq. 2 with the model in Eq. 1, it follows that the expression data **Y***_G,C_* conditioned upon *ω*_*T,C*_ obeys:

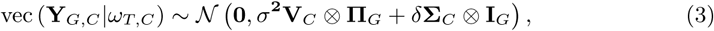

with 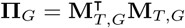, where ^T^ is the transpose. Hence, the covariance between genes is driven by the similarity among motif counts **M***_T,G_*. **Σ***_C_* is the covariance of noise between conditions and *δ***I***_G_* is the covariance matrix between genes. Thus, we assume independence between genes in the noise term. The posterior mean of *ω*_*T,C*_ given vec (**Y***_G,C_*), denoted 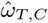, then reads:

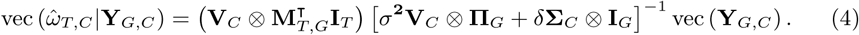

This model is explained in more detail in Supplement A.

#### Special Case: Ridge Regression

Ridge Regression, also known as Tihkonov Regularization, prevents over-fitting in a linear regression by using an *ℒ*^2^-regularization on the estimated parameters. For more details on *ℒ*^2^ -regularization and Ridge Regression, we refer the reader to Ng (2004). Note that in a Ridge Regression the samples are assumed to be isotropic, i.e. independent and identically distributed. The covariances in the model, **V***_C_* and **Σ***_C_*, therefore reduce to identity matrices which only differ in the constant with which they are multiplied:

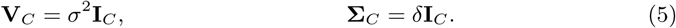

With the above model we formulated a Bayesian linear mixed model to explain the motif influence on expression data **Y***_G,C_* based on motif scores. The Bayesian Linear Mixed Model provides a relaxation of the so far used assumption of independence in Ridge Regression. It allows for modeling dependent structures between conditions and in the noise, which can increase the power of the model, as shown in the Results section. In the following, we will refer to the model as Bayesian Linear Mixed Model when allowing for correlation between samples and noise, i.e. the covariance matrices **V***_C_* and **Σ***_C_* are not restricted except for being symmetric positive definite matrices. We refer to the model as Ridge Regression when we specifically assume independence between conditions and noise, i.e. **V***_C_* = *σ*^2^**I***_C_*, and **Σ***_C_* = *δ***I***_C_*,

### Implementation and model testing

The project implementation (https://github.com/Sim19/SimGEXPwMotifs) is provided in the programming language Python (Oliphant, 2007; Millman and Aivazis, 2011), where we make use of the packages `pandas` (McKinney, 2010), `matplotlib` (Hunter, 2007) and `numpy` (Oliphant, 2006). We run the optimization of Eq. 3 to compute Eq. 4 with the Python package `limix` (Lippert *et al*., 2015, 2014a; Casale *et al*., 2015) and use the module `VarianceDecomposition`, which we use with the parameter settings for restrictions on **V***_C_* to “freeform”, which models concurrently **Σ***_C_* to be of “random” shape. These parameter settings only restrict **V***_C_* and **Σ***_C_* to be positive-definite matrices, with 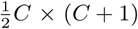 parameters to be estimated. With these settings, there are no restrictions on the rank of the matrix, i.e. the degree of correlation among samples. For the optimization start, **V***_C_* and **Σ***_C_* are both set to be the estimated covariance matrix of 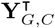, each divided by half. The optimization is based on the *L-BFGS* algorithm to minimize the log likelihood with an *ℒ*^2^-regularization along the non-diagonal elements of the covariance matrix **Σ***_C_*, which is also known as isotropic Gaussian prior with zero mean. The implementation makes use of the reduction of computational complexity by using the Kronecker product notation and its identities for the case of matrix variate data, which is highly efficient (Lippert *et al*., 2014a; Casale *et al*., 2015). For more detailed information about Linear Mixed Models and its implementation in `limix`, refer to Lippert *et al.* (2014b). We run Ridge Regression in the simulation study with the Python implementation in the package `sklearn.linear model.RidgeCV` (Pedregosa *et al*., 2011). For the data application, we make use of the `limix.VarianceDecomposition` implementation with the setting “freeform” for **V***_C_* and “random” noise for **Σ***_C_* for fitting the Bayesian Linear Mixed Model. For the computation of Ridge Regression, we restrict **V***_C_* and **Σ***_C_* to be identity matrices. Both computations (`limix.VarianceDecomposition` with restrictions for **V***_C_* and **Σ***_C_* to identity matrix and `sklearn.linear model.RidgeCV`) yield the same results (see Fig. 1 in Supplement C).

**Figure 1:**
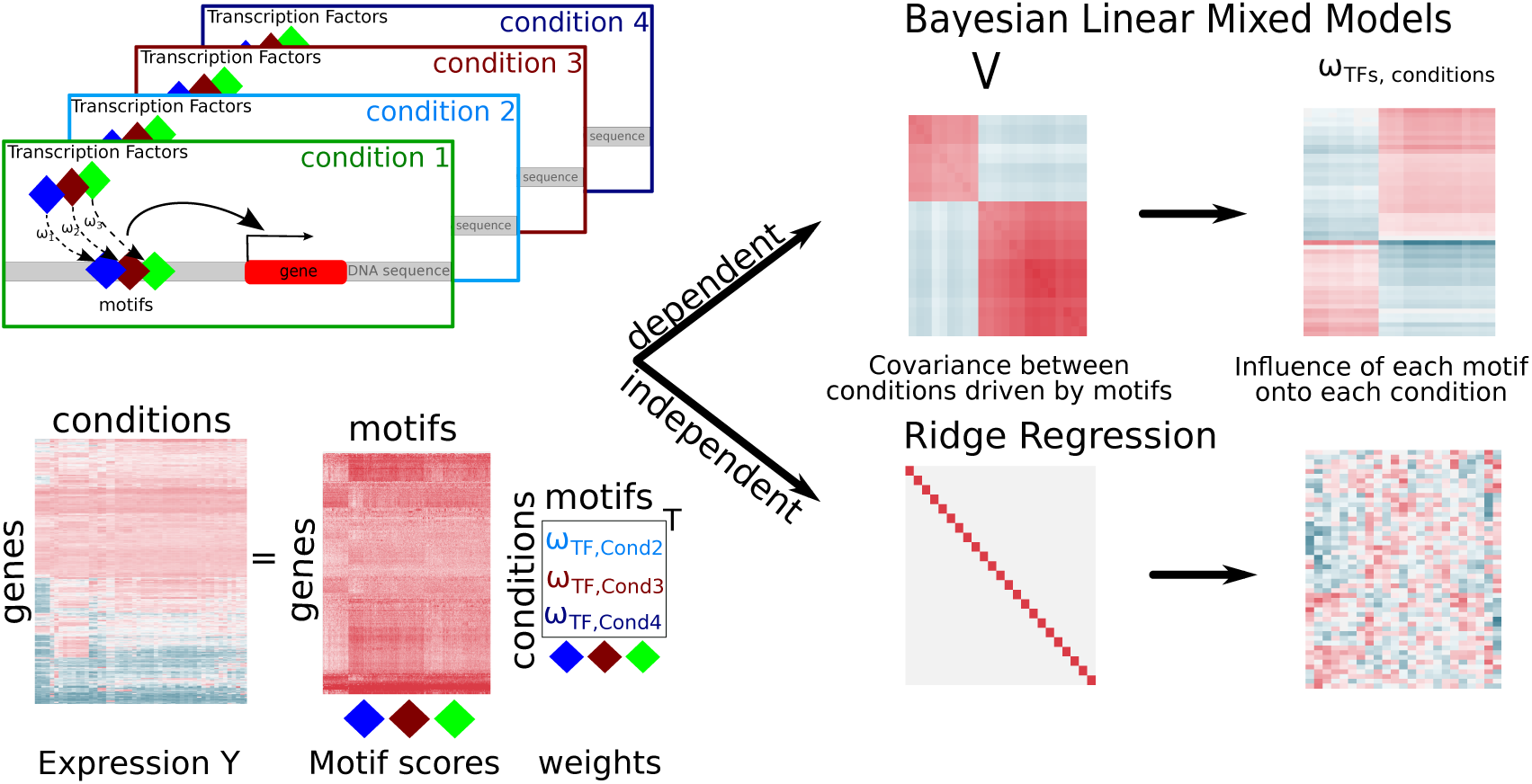
Project Overview.

For the visualizations in this article, we work with the plotting facility from `pandas` and `seaborn` (Waskom *et al*., 2017). For boxplots and other results from the simulation study, we make use of the `R-ggplot2` package (Wickham, 2016) and `cowplot` (Wilke, 2018). The visualization of clustered data is done with python’s `seaborn.clustermap()` using the “complete” method for the dendogram computation on Euclidean distance.

### Simulating data

For the simulation study we generate data based on the model introduced in Eq. 1 - Eq. 4, given a covariance matrix **V***_C_*. The prior weight 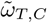 and the expression signal **Y***_G,C_* are generated according to Eq. 2 and Eq. 1. The expression data **Y***_G,C_* from Eq. 1 is then used to estimate **V***_C_* and **Σ***_C_*. Based on these computations, the posterior motif influence 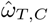 is then computed and compared to the simulated motif influence 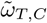 with a Pearson correlation (Pearson, 1895) over all conditions. As gene set we use the 978 landmark genes from the LINCS project (Koleti *et al*., 2018). In a secondary simulation we increase the size of the gene set to 5000 genes, which originate from an analysis of the most variational genes across all samples from the GTEx project (Genotype Tissue Expression, https://gtexportal.org/home/). We generate data for *C* = {10, 30, 50, 70, 100, 120} conditions.

### Covariance types of V*_C_*

For the generation of simulated motif influence 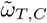 (Eq. 2), we need to give a covariance matrix *σ*^**2**^**V***_C_* ⊗ **I***_T_*. As the covariance along TFs is modeled to be isotropic (independent and identically distributed), one can generate *ω*_*t,C*_ randomly *T* times with *ω*_*t,C*_ *∼ 𝒩* (0, *σ*^**2**^**V***_C_*). For the covariance between the weights *ω*_*T,C*_ in Eq. 2 and hence expression data **Y***_G,C_* in Eq. 3, we consider four types of covariance matrices for **V***_C_*, according to assumptions made about the correlation between samples: (i) Isotropic V: same number on the diagonal, all off-diagonal elements set to zero. Samples drawn from such a covariance matrix are independent. This matches the implicit assumption of the Ridge Regression. We will refer to this as “V: independent”. (ii) Full matrix V with all positive off-diagonal elements. Samples drawn from such a covariance matrix are positively correlated, without any specific (block) structure. This is a clear mismatch with the assumptions underlying Ridge Regression, and results are expected to favor the (full) Bayesian Linear Mixed Model. We will refer to this as “**V***_C_* : correlated (no groups)”. (iii) Block structure with many blocks, each modeling a group of biological replicates. Samples within each block are highly correlated, small correlation between the blocks. Blocks vary in size. Average block size is two. We will refer to this as “**V***_C_* : correlated (many groups)”. (iv) Block structure similar to (iii), but now with just two blocks. We will refer to this as “**V***_C_* : correlated (two groups)”.

For the generation of the correlated covariance matrices with groups, we provide pseudocode in Supplement. An exemplary visualization of a covariance matrix with samples that correlate in many groups is given in Fig. 2A and B, left panel.

**Figure 2:**
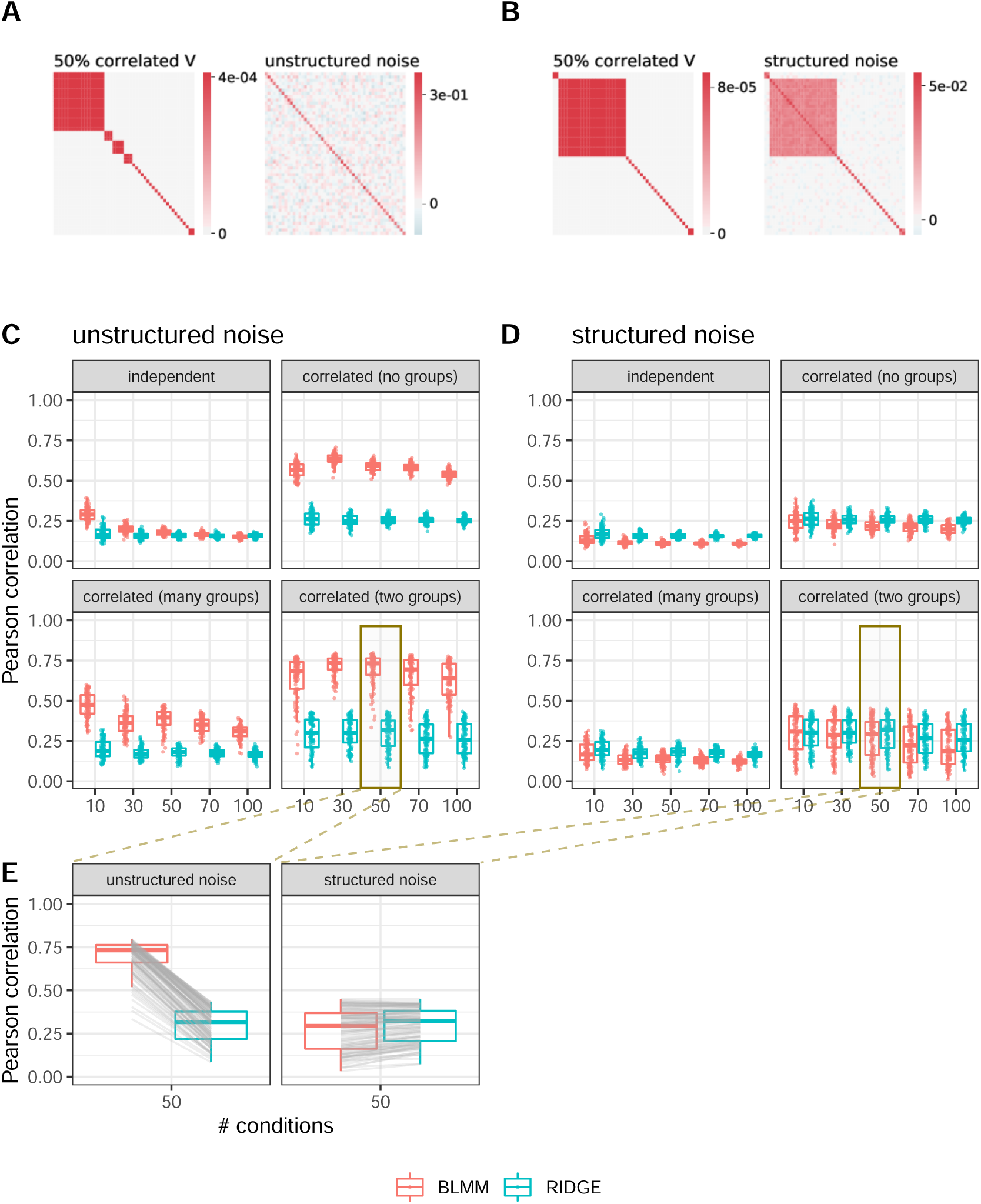
Simulation Study. Data is generated over *G* = 978 informative genes and *T* = 623 motif scores and 100 repetitions, with either unstructured noise **Σ***_C_* (A,C) or structured noise **Σ***_C_* (B,D). (A,B): exemplary covariance matrix between samples, **V***_C_*, and noise matrix **Σ***_C_* being unstructured (A) or structured (B). Data is shown for *C* = 50 samples with a covariance between conditions with *k* = 25 blocks, i.e. 25 sample groups in which samples are completely correlated. (C) and (D): The Pearson correlation values between generated and estimated motif-condition-weights are shown for different degrees of correlation between samples. The values are depicted per method used to predict the motif-condition-weights: the Bayesian Linear Mixed Model (BLMM, in red), which allows for dependence between samples, and the Ridge Regression (RIDGE, in blue), which assumes independences between samples. Data is generated with the following covariances between samples: (i) independence between samples, **V***_C_* = **I***_C_*, (ii) unrestricted correlation between samples, (iii) correlated data with many sample groups, and (iv) highly correlated samples, assuming samples originate from two biologically different samples, Results are shown based on data generated with unstructured noise ((C), see (A)), or with structured noise ((D), see (B)). (E): Exemplary comparison of Pearson correlation values between simulated and predicted posterior motif influence 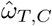. The data is generated with highly correlated samples, modeling two groups of biological replicates. Corresponding replicates are combined by a gray line. The data in the left panel is generated with unstructured noise **Σ***_C_* (see (A)) and in the right panel with structured noise (see (B)).

### Noise Σ*_C_* - unstructured and structured

To generate the gene expression data **Y***_G,C_*|*ω*_*T,C*_, we compute the signal as the product of the motif scores **M***_T,G_* (explained hereafter) and their weights *ω*_*T,C*_ (explained previously). Due to the randomness in the signal that is not explained by motifs, we add some random noise which is drawn from a normal distribution with covariance *δ***Σ***_C_* ⊗ **I***_G_*. We generate **Σ***_C_* in two different ways: (i) we assume no particular structure, **Σ**_*C*,random_, which is a matrix filled with values drawn from a standard normal distribution, multiplied with itself. This is the Wishart distribution with *C* degrees of freedom. Or (ii) we add the same covariance matrix **V***_C_* that is used to model the correlations between the conditions to **Σ**_*C*,random_ from (i): 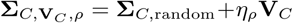. The noise matrices are then normalized by their trace. The detailed description of the generation of the noise matrices as well as the control of structuredness in it is given in Supplement. For the simulation study depicted in Fig. 2 and discussed in the Results section we generate the structured noise with *ρ* = 0.7. A visualization of both types of noise matrices is given in Fig. 2A and B, both in the respective right panel.

#### Signal-to-noise ratio

Previous research has shown that roughly 10 *-* 20% of the signal of gene expression can be explained by motif influence in the promoter region (Balwierz *et al*., 2014). We therefore generate the data in such a way, that 20% of the signal in expression data **Y***_G,C_* is due to motifs, and the rest unexplainable noise. We achieve this by adjusting the parameter *σ*^**2**^ and *δ*. We fix *δ* and determine *σ*^**2**^ by bisection such that it explains 0.2 of the variance coefficient (Eq. 1). More details can be found in Supplement.

### Motif scores

For the computation of the transcription factor motif scores **M***_T,G_* we use log-odds scores based on the positional frequency matrices, which are computed with the software `GimmeMotifs`, v13 (van Heeringen and Veenstra, 2011; Bruse and van Heeringen, 2018). We make use of the database `gimme.vertebrate.v3.1`, included with `GimmeMotifs`. We assume the promoter region to be 400bp upstream and 100bp downstream of the Transcription Start Site of the given gene sets. We take the Transcription Start Site from the GENCODE database (version 26) (Frankish *et al*., 2019) and generate a `bed` file with the promoter regions with `BEDTools, v2.17.0` (Quinlan and Hall, 2010) and the subcom-mand `slop`. In general, we filter for genes that are known to be protein coding and on chromosomes 1-22 and X.

### Cross-Validation

In the Results section, we compare the performance of Bayesian Linear Mixed Model and Ridge Regression on two real-world datasets. To assess the performance of both model assumptions, we make use of a ten-fold cross-validation by creating ten random subsets across the genes. Each of these ten subsets is used as a test dataset, while the model is trained on the union of the remaining nine subsets. As there is no knowledge about the motif influence 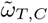, we compute the expression 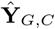 with the predicted posterior motif influence 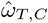 (Eq. 1) and compare the Pearson correlation between predicted expression 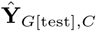 on the test set and the original expression **Y**_*G*[test],*C*_ of the test set.

### Experimental Data

For the application analysis presented in the Results section, we make use of two experimental datasets.

#### H3K27ac ChIP-sequencing signal at hematopoietic enhancers

We use experimental enhancer activity data from the human hematopoietic lineage (Martens and Stunnenberg, 2013; Bruse and van Heeringen, 2018). This dataset is based on 193 ChIP-seq experiments in 33 hematopoietic cell types using an antibody specific for histone H3 acetylated at Lysine 27 (H3K27ac). The H3K27ac histone modification is deposited by the histone acetyltransferase p300 (EP300) and is associated with enhancer activity (Creyghton *et al*., 2010). ChIP-seq reads were counted in 2kb regions centered at accessible regions, log-transformed and normalized using scaling by Z-score transformation. The peaks, or accessible regions, represent putative enhancers. We subset the peaks to the most variable 1000 peaks over all samples. We subset for all replicates of the cell types “monocytes” and “T-cells”, which are 23 samples in total. Different samples of the same cell type represent different donors.

#### Human tissue gene expression data (GTEx)

Second, we make use of gene expression data from human tissues. The data is from the Genotype Tissue Expression (GTEx) database (Lonsdale *et al*., 2013a), and is available on https://gtexportal.org/home/. It is RNA-seq data from many different tissues. The data was downloaded with project number *SRP* 012682 with the R-package `recount`, v.1.63 (Lonsdale *et al*., 2013b; Collado-Torres *et al*., 2017b,a; Ellis *et al*., 2018). We scale the raw counts by the total coverage of the sample (function `scale counts()`, setting “by=auc”) and keep entries with at least 5 counts. We transform the data with the `DESeq2`-package, v.1.20.0 and use the variance stabilizing transformations (Tibshirani, 1988; Huber *et al*., 2003; Anders and Huber, 2010), implemented in the package function vst(), with `blind` transformations to the sample information.

We selected the 5, 000 most variable genes over all samples of the entire GTEx experiment. We then choose 75 random samples, of which there are 35 different tissues from 21 different organs.

As we model the normalized expression data **Y***_G,C_* (see Eq. 1), we subtract the mean along genes and along conditions from the expression data. For the analysis, we further normalize the motif score matrix.

## Results

In this section, we apply the model, that we introduced in detail in Eq. 1-Eq. 4, to simulated data and to two real-world datasets. We compare the two assumptions about the shape of covariance, as discussed before: the novel allowance of dependence (Bayesian Linear Mixed Model) to the restriction of independence, that was applied so far (Ridge Regression).

### Simulation

To quantify the differences in the model when allowing for dependence between conditions, instead of assuming independence, we simulate data according to Eq. 1. We generate different datasets for *G* = 978 genes, for *C* = *{*10, 30, 50, 70, 100, 120*}* samples, and *T* = 623 motifs. We vary the degree of correlation between the samples, expressed in the covariance matrices **V***_C_* and **Σ***_C_*. We also vary the influence of these covariance matrices on the signal: As covariance matrix **V***_C_* we generate (i) an identity matrix, (ii) a matrix with unrestricted correlation between samples, (iii) a lower rank matrix with 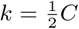 assumed groups of biological replicates and (iv) a lower rank matrix with *k* = 2 groups of biological replicates. As noise matrix **Σ***_C_* we first generate unstructured or random noise. In a second step, we generate the data with a structured noise matrix **Σ***_C_*. For every parameter set we generate 100 replicates. We compare the performance of the two assumptions on the shape of covariance with a Pearson correlation score. The correlation is computed between the generated simulated motif influence 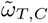 and the estimated posterior motif influence 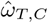. The higher the correlation values the better the performance of the model. In Fig. 1, we confirm that the Bayesian Linear Mixed Model with fitting independent samples and independent noise is equal to Ridge Regression.

We summarize the simulation study visually in Fig. 2. We separate the simulation study first on data generated with unstructured noise (exemplary visualization of data in Fig. 2A, Pearson correlation values in Fig. 2C) and second on data with structured noise (exemplary data shown in Fig. 2B, model performances shown in Fig. 2D). Summarizing, we compare the performance of both models on the two types of data (unstructured versus structured noise **Σ***_C_*) in Fig. 2E. We show an example of the model performance for data generated with a covariance matrix **V***_C_* that models all samples to originate from many blocks on unstructured (left) and structured noise (right). In Fig. 3, we show on the example of *C* = 50 samples that a higher dimensional dataset, generated with *G* = 5000 genes, exhibits the same performance as for *G* = 978 genes.

**Figure 3:**
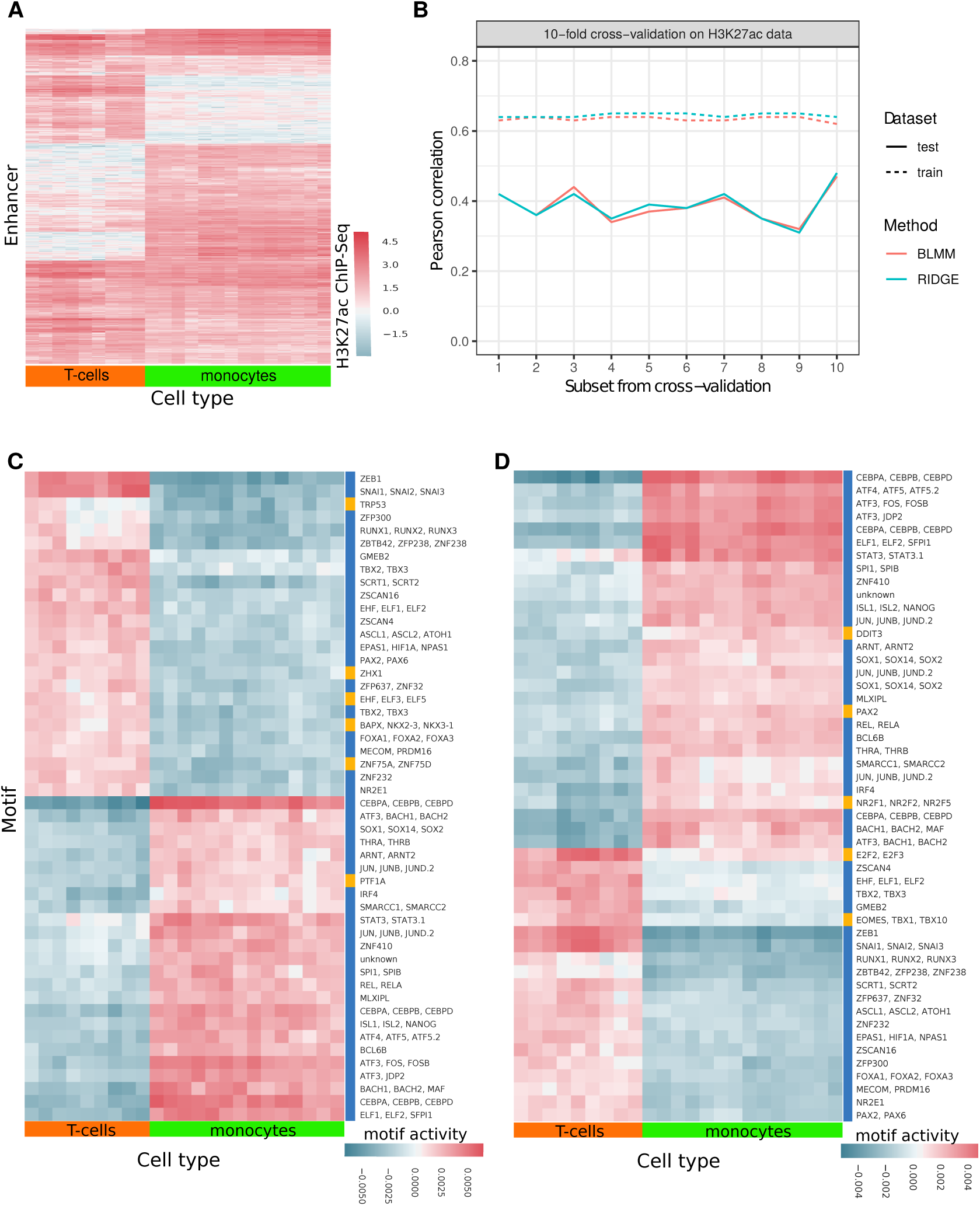
Data application on H3K27ac data. (A): Pattern of H3K27ac ChIP-seq signal in putative enhancers over two hematopoietic cell types, T cells and monocytes. (B): Pearson correlation between training set and predicted ChIP-seq signal on training set and between test set and predicted test ChIP-seq signal of a ten-fold cross-validation. (C) and (D): Motif weights for 50 enhancers across the two cell types, assuming dependence (C) or independence (D) between the conditions. Enhancers are chosen based on largest difference in mean H3K27ac ChIP-seq signal per tissue group. Common motifs of Bayesian Linear Mixed Model (C) and Ridge Regression (D) are depicted in blue, others are depicted in yellow.

### Unstructured noise Σ*_C_*

Given the covariance **V***_C_*, we generate a gene expression signal **Y***_G,C_* according to Eq. 1 with random noise **Σ**_*C*,random_ (see Fig. 2A). In Fig. 2C, we compare the performance of Bayesian Linear Mixed Model, which allows for dependencies between samples, to Ridge Regression, which assumes independence. We depict the Pearson correlation values between estimated and simulated motif influence *ω*_*T,C*_ for 100 datasets per boxplot. In every panel, we compare both model assumptions, (i) dependence (in red, labeled BLMM) and (ii) independence (in blue, labeled RIDGE). We present the results separated by the degree of correlation in **V***_C_*, which was used to generate the data. Allowing for dependencies between conditions leads to a better prediction performance, especially for correlated data (Fig. 2C, E), independent of the sample size *C*. For uncorrelated data (**V***_C_* = *σ*^2^**I***_C_*), the allowance for dependency yields a slightly better performance for small sample sizes, *C ∈ {*10, 30, 50*}* (Fig. 2C, upper left panel). For higher sample sizes, the performances are equal. In Fig. 2E, left panel, we show explicitly that the Pearson correlation values resulting from the dependence assumption (BLMM, in red) are always higher than those from the independence assumption (RIDGE, in blue). The exemplary visualization shows the correlation between estimated and simulated motif influence 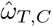, which was generated with a covariance matrix **V***_C_* of rank 2, i.e. *k* = 2 correlated groups of samples over *C* = 50 conditions. We run both model assumptions on the same datasets. The correlation values from the same datasets, depicted per model assumption, are connected with gray lines.

#### Structured noise Σ*_C_*

For the generation of the gene expression signal with structured noise, we add the structure of correlation between conditions to the noise (see Fig. 2B). This is motivated by the fact that we can explain roughly 20% of the signal in expression data **Y***_G,C_* data by motif influence, but not the remaining 80% of the signal. This remaining signal is often similar to the covariance between samples. In Fig. 2D, the Pearson correlation values are depicted that are computed between simulated motif influence 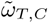 and posterior motif influence 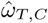. They result from applying the dependence (BLMM, in red) and independence assumption (RIDGE, in blue) to data that is generated with such a structured noise at a degree of 0.7. The results are again shown for differently correlated datasets between conditions **V***_C_*, analogous to the previous section. With that structure in the noise **Σ***_C_*, both methods perform equally with low correlation values. As shown in Fig. 2E, right panel, the Bayesian Linear Mixed Model performs equally or slightly worse than Ridge Regression.

Indeed, when applying a degree of structure in the noise **Σ***_C_* (explained in detail in Eq. 17 - Eq. 18 in Supplement) with varying *ρ* between zero and one in a step size of 0.1, there is loss of performance of the Bayesian Linear Mixed Model for an increasing structured signal in the noise (see Fig. 2). In contrast, the correlation values resulting from Ridge Regression applied to this data with structured noise are very similar, if not equal, to those from data that was generated with unstructured noise (Fig. 2C). Both model assumptions perform approximately equally when applied to data with structured noise. For an exemplary visualization we depict the correlation values per sample in Fig. 2E, right panel, with lines connecting the two results from the two model assumptions, which were applied to the same dataset. The reason for this performance loss is the computation of posterior motif influence 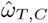 (Eq. 4). If **Σ***_C_* takes a structure that is too similar to **V***_C_*, these two terms can be summarized in the covariance of vec (**Y***_G,C_*|*ω*_*T,C*_) in Eq. 3. This covariance returns as inverse in Eq. 4 and therefore **V***_C_* and **Σ***_C_* cancel out with **V***_C_* from the motif-dependent term 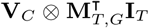 (see Eq. 4). We give more mathematical details in the discussion.

### Application

We compare the Bayesian Linear Mixed Model and the Ridge Regression on two real-world datasets. First, we apply the method to a ChIP-seq dataset of the histone modification H3K27ac in the human hematopoietic lineage. This histone mark correlates with enhancer activity (Creyghton *et al*., 2010). We refer to this dataset as H3K27ac. Second, we use RNA-seq gene expression data from the GTEx consortium. It is known from RNA-seq data, that there is generally a high contribution of “technical” noise, such as measurement noise that is introduced purely by the experiment (laboratory (Irizarry *et al*., 2005; Shi *et al*., 2006) and batch (Leek *et al*., 2010)) and by biological variation (Hansen *et al*., 2011). Hence we expect a significant contribution of such “technical” noise to the signal. We refer to this RNA-seq dataset as GTEx.

For both datasets, we compare the dependence and independence assumption (Bayesian Linear Mixed Model and Ridge Regression) by means of a cross-validation.

#### Acetylation data H3K27ac

We split the acetylation dataset H3K27ac into two sample groups, originating from two different cell types, (i) monocytes from the myeloid lineage and (ii) T-cells from the lymphoid lineage. Within these two groups the ChIP-seq signal **Y***_G,C_* is highly correlated as the samples represent the same cell types from different donors. The results from the application of the Bayesian Linear Mixed Model and the Ridge Regression to the H3K27ac dataset are summarized in Fig. 3.

To assess and compare the model performances, we run a ten-fold cross-validation. Per run we compute the Pearson correlation between measured and predicted ChIP-seq signal **Y***_G,C_* of the test and training set. The correlation values from the cross-validation are shown in Fig. 3B. The performance of the test set is depicted to gain insight into the prediction performance of each model, the performance of the training set to check for over-fitting. Both model assumptions yield very similar performances. Hence, on the basis of cross-validation, no clear statement about superiority of one model assumption to another can be made. A visualization of the estimated **V***_C_* and **Σ***_C_*, depicted in Fig. 4 - Fig. 7, emphasizes again the great difference between the two models and their model fits. While for Ridge Regression both matrices are identity matrices with a scaling factor, both covariance matrices from the more flexible Bayesian Linear Mixed Model (Fig. 6 and Fig. 7) exhibit strong correlation structures. These two latter matrices are very similar. Clustering the covariance matrices yields two clearly separated blocks along the diagonal, showing a strong correlation among the samples within the cell types, and independence, or even a slight anti-correlation, between the two cell types. This pattern follows the partitioning of the samples into biological replicates for the two different cell types.

**Figure 4:**
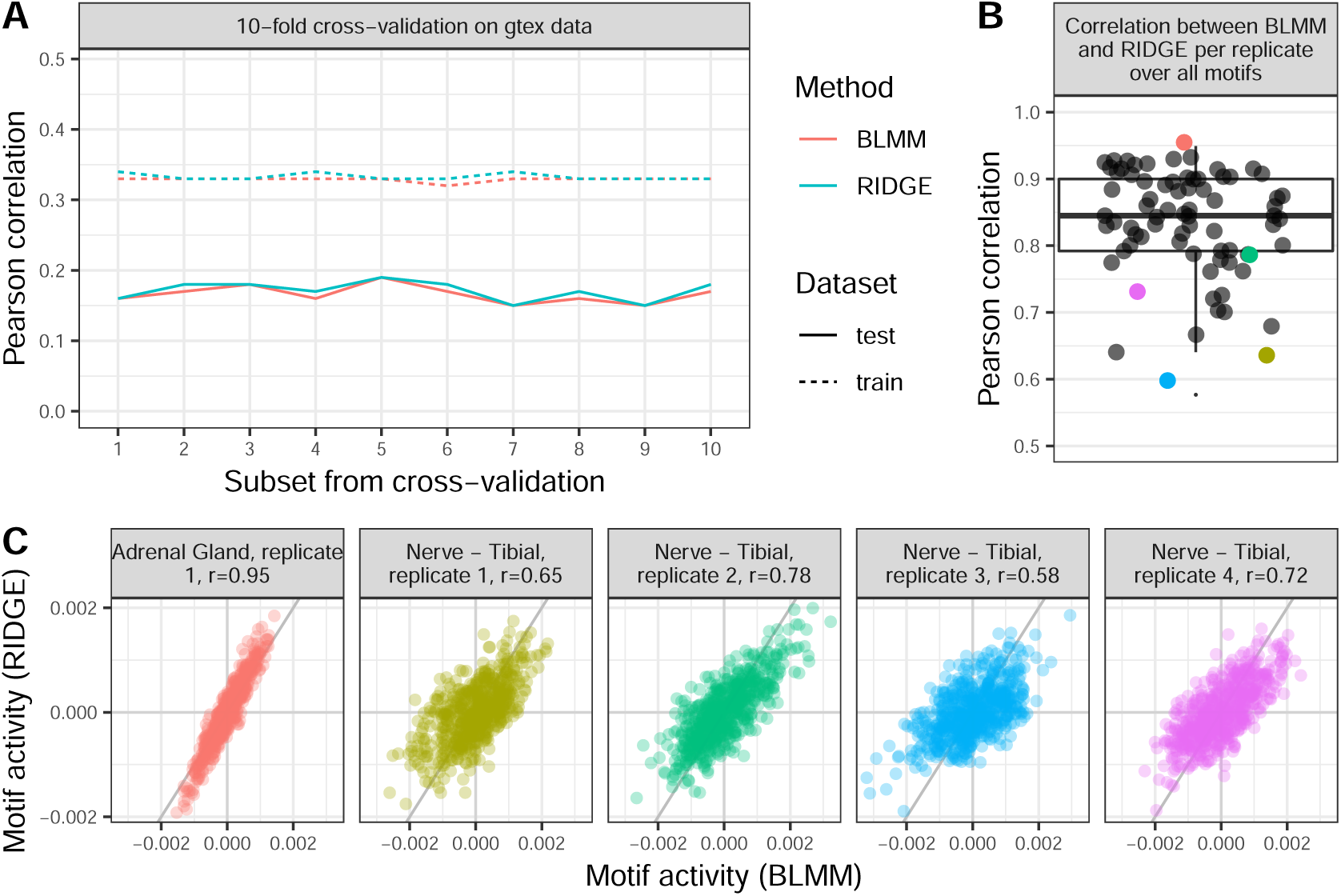
Data application on GTEx data. (A): Pearson correlation values between training set and predicted gene expression of training set and between the test set of the gene expression data and the predicted gene expression of ten-fold cross-validation for Bayesian Linear Mixed Model (BLMM, in red) and Ridge Regression (RIDGE, in blue). (B): Pearson correlation of estimated motif scores on the basis of the Ridge Regression and the Bayesian Linear Mixed Model. Per correlation value, all motif scores are taken per replicate. Examples shown in (C) are colored according to respective color. (C): Exemplary scatter plots of tissue replicates on which the Pearson correlation for the posterior motif influence 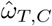 is high or low between the two methods, Bayesian Linear Mixed Model and Ridge Regression, as colored in (B). The values along the x-axis result from the Bayesian Linear Mixed Model, assuming dependence betweem samples, and on the y-axis from assuming independence (Ridge Regression).

Looking in more detail into the importance of the posterior motif influence 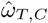, we compare the motifs by ranks. They are ordered based on the difference of the motif’s mean influence onto each class, i.e.

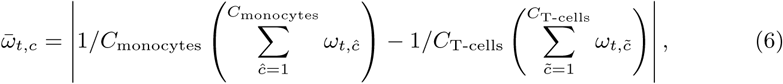

with *C*_monocytes_ + *C*_T-cells_ = *C*. They are sorted in decreasing order, i.e. the higher the mean difference, the higher the rank of a motif, where rank 1 is the most important motif and rank 623 the least important. For Fig. 3C and D, we chose the top 50 ranking motifs, respectively. We color motifs that are shared for both methods in blue and different motifs in yellow. All in all, 40 motifs are shared among the 50 chosen. The posterior weights, depicted in Fig. 3C and D, clearly separate the motif influences based on the underlying cell types for both models. Despite the clear differences in model estimates, the performances in predictive power are very similar. We hypothesize that the similarity of the performances results from the similarity between estimated covariance matrix **V***_C_* and noise **Σ***_C_* in the Bayesian Linear Mixed Model, which leads to canceling out the covariance structure in the posterior. We elaborate on this phenomenon in the discussion.

#### RNA-seq data GTEx

We further compare the performance of both model assumptions on a human tissue-specific RNA-seq dataset from GTEx. In comparison to the H3K27ac dataset discussed in the previous section, RNA-seq is known to exhibit more “technical” noise, i.e. noise that is added to the signal purely by conducting the experiment. Hence the expected noise structure should be better separable from the motif signal.

For the *G* = 5000 most variational genes across the entire GTEx dataset, we apply the Bayesian Linear Mixed Model and the Ridge Regression on *C* = 75 randomly chosen samples. Among those, there are several biological replicates, resulting in 35 different tissue types across 21 organs. The results of the GTEx dataset are summarized in Fig. 4.

Analogous to the analysis on the H3K27ac dataset, we conduct a ten-fold cross-validation. The Pearson correlation values of expression data **Y***_G_*,_*C*_ are very similar for both model assumptions (see Fig. 4A). The Pearson correlation of all motif scores between the two model assumptions per tissue is high with the median at 0.83, the first quantile at 0.79 and the third quantile at 0.90 (see Fig. 4B). In Fig. 4C we highlight examples of high (left) and low (remaining four panels on the right, same replicate) Pearson correlation values between the estimated motif weights from the two models (colored accordingly).

When we compare the estimated covariance matrices, there are strong differences in **V***_C_* and **Σ***_C_* (Fig. 9 - Fig. 12). Independent of the model assumptions, there is a difference of 10^4^ in order of magnitude of the signal assigned to the covariance between condition and the estimated noise.

We compute the inter-quantile range of the posterior weights of all *T* = 623 motifs per tissue, to investigate the posterior motif influence 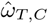 in more detail. We summarize the posterior weights over tissues with replicates with the median. We then filter those motifs that lie outside the range of 2.5 times the inter-quantile range above and below the median. Combining the two sets from Bayesian Linear Mixed Model and Ridge Regression results in 56 motifs, of which 50 are in the intersection of the two sets.

The clustering of posterior motif influence 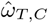 results in comparable clusters (Fig. 8), with the clustering on tissues from Ridge Regression being slightly better due to a clearer clustering of replicates.

The overall correlation between the posterior motif influence 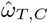 per tissue of both methods is very high (median *≥* 0.95, Fig. 5A), which underlines the strong similarity between the two models (see Fig. 13 for the five highest correlated motifs across all tissues and Fig. 14 for the five lowest correlated motifs). The assumption of dependence between samples (BLMM) results in higher variation of posterior weights (see Fig. 15). Among the 56 chosen motifs, there are a few cases, where one model assumption yields a weight value of around zero and the other is significantly different from zero (see Fig. 5B). Out of those four examples, we found evidence that the TFs are known to play a major role in the descriptive tissue: *FOXO1, FOXO3* and *FOXO4* in liver (Tikhanovich *et al*., 2013), *RUNX1-3* in blood (Sood *et al*., 2017; Okuda *et al*., 2001; Qiao *et al*., 2006) and in B-cell lymphocytes (Gunnell *et al*., 2016). We found no explicit relevance for *YY1* in skin fibroblasts. The *YY1* TF is known to be involved in the repression and activation of a diverse number of promoters (Taguchi *et al*., 2011; Bollag and Bollag, 2011; National Center for Biotechnology Information (US), 2019). It is relatively highly expressed in skin cells and fibroblasts, which we find from gene expression profiles from the Protein Atlas (Uhlen *et al*., 2017), data available from `v18.1proteinatlas.org`. This could be biologically relevant, but no specific function is known. Hence, despite similar performances we find evidence of motif influences found by the Bayesian Linear Mixed Model, but not by Ridge Regression. The same holds vice-versa for EBV-transformed lymphocytes, for which Ridge Regression predicts an influence of the motif to which the TFs *RUNX1, RUNX2, RUNX3* bind, whilst the Bayesian Linear Mixed Model does not. This finding is concordant with the observation that the Epstein-Barr virus (EBV) TF EBNA-2 induces RUNX3 expression in EBV-transformed lymphocytes (Spender *et al*., 2002).

**Figure 5:**
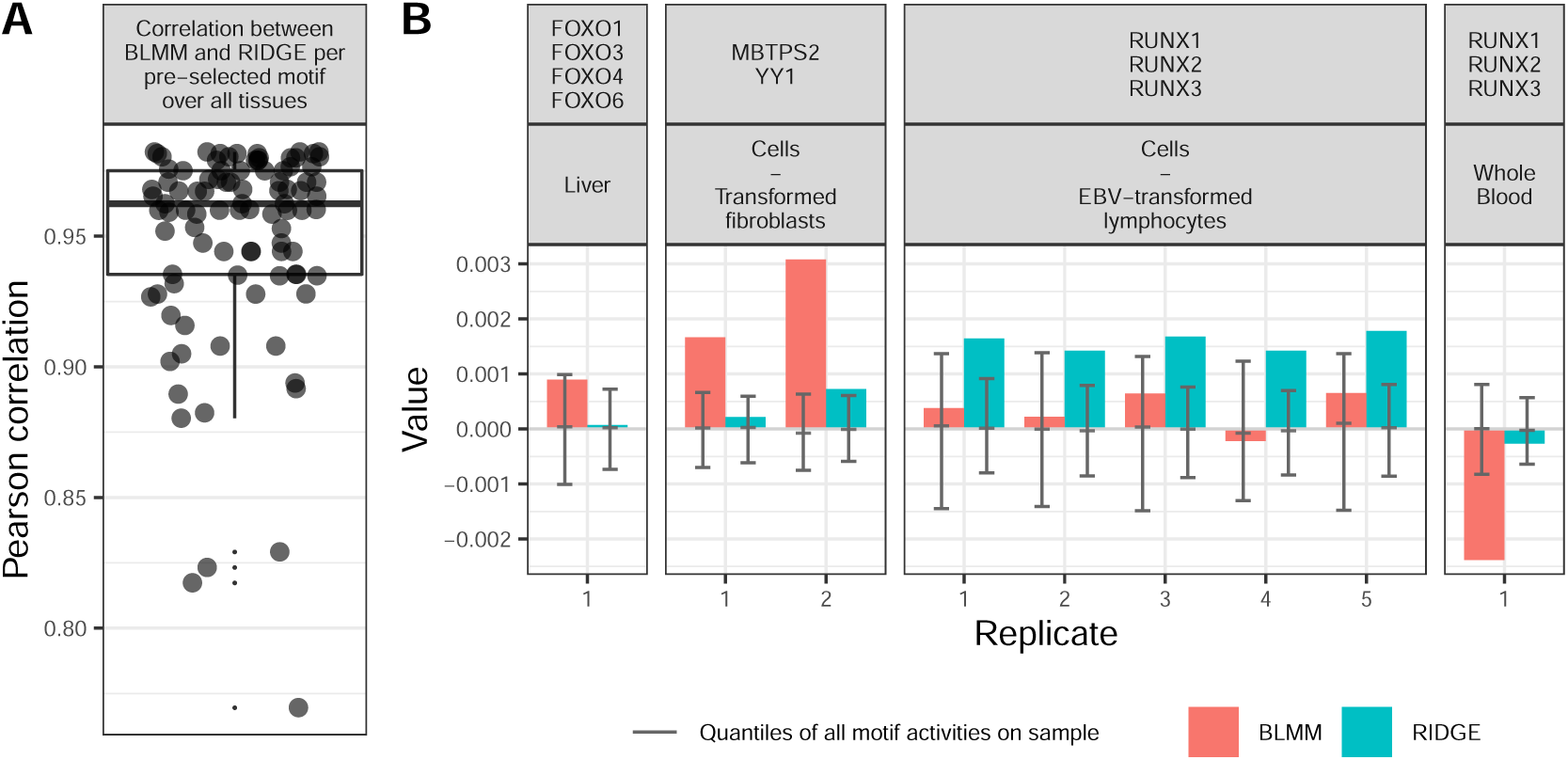
IQR-study on GTEx motif-weight. (A): Pearson correlation values between the predicted posterior motif influence 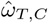 from the Bayesian Linear Mixed Model and the Ridge Regression of 56 selected motif values over all tissues. (B): extreme cases of different motif scores per method. Each box shows the predicted motif scores per sample, separated by model assumptions used to predict the scores (dependence, denoted BLMM, colored in red, and indepdence between samples, named RIDGE, colored in blue). Per sample, the first, second and third quantile (as in a boxplot) of the overall motif activity of all motifs on that sample are depicted in gray.

## Discussion

We observe a clear superiority of the Bayesian Linear Mixed Model over Ridge Regression in the Results section, if the entire signal can be explained by motif influence and no or less comparable structure between samples can be found in the noise. With the simulation study, we observe a decrease in performance for the Bayesian Linear Mixed Model for an increasingly structured signal in the noise (see Fig. 2). When applying the two model assumptions to two real-world datasets, we see no favorable performance on the basis of a cross validation. Only a more detailed investigation of motif importance reveals some differences, which are a lot stronger for the GTEx RNA-seq dataset, as more “technical” noise is present than in the acetylation dataset. The origin of noise in our model has two different sources: the “technical” noise, which includes noise introduced (i) in the lab (different technicians, different days of experiment conducted, pipeting error, different kits, etc.), (ii) by the machine (batch effect, sequencing, lane-to-lane variability, etc.), and (iii) through biological variability due to gene expression being a stochastic process (Elowitz *et al*., 2002). The other source of noise originates from the model, that explains the expression signal uniquely as a linear combination of motif scores. Other sources that contribute to the signal are not modeled here, hence end up in the noise. And it is this large contribution of noise to the signal, which is no “technical” noise, that causes the loss of performance of the dependence assumption in the covariance. Mathematically, **V***_C_* cancels out of the computation of the posterior (Eq. 4) if the covariance between conditions **V***_C_* and the noise term **Σ***_C_* are comparable. This is the reason to why the Bayesian Linear Mixed Model does not perform better than the Ridge Regression in this case. If the noise takes a form similar to the correlation between conditions, say:

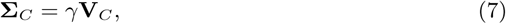

with some constant scaling *γ*, then Eq. 4 takes the following form:

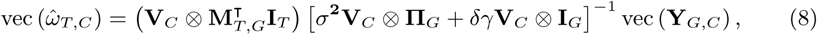

where **V***_C_* cancels out in the equation:

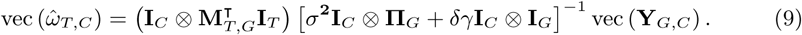

Hence, the correlation structure between samples plays no role in the determination of posterior motif influence 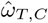. One can therefore conclude, that the entire formulation of the Bayesian Linear Mixed Model we proposed is equivalent to Ridge Regression if **V***_C_* and **Σ***_C_* are the same up to a scaling factor. Hence, the Bayesian Linear Mixed Model is only to favor over Ridge Regression if there is less noise from the signal than from the “technical” noise in the data.

With the provided Bayesian formulation of Ridge Regression, we extended the model by allowing for correlation between samples and for correlated noise. We used this extension here for the investigation of motif influence on gene expression at the promoter region. Our approach could be extended by taking into account motif information of enhancers. Furthermore, it can be combined with other possible influential sources, such as chromatin accessibility and chromatin interaction maps determined by chromosome conformation capture techniques such as Hi-C (Lieberman-Aiden *et al*., 2009). Also, the number of motifs used could be extended to a more complete set (Lambert *et al*., 2019).

The relaxation of the assumption of independence between samples in the model allows for the consideration of correlation and hence to better control the breakdown of measured signal into different sources. There are many applications in the field of molecular biology, where Ridge Regression has proved to be successful. The limix package itself was mainly developed to investigate the influence of SNPs on phenotype prediction (Stegle and Lawrence, 2012; Rakitsch *et al*., 2013; Casale *et al*., 2015). With an increase of computational power and better implementations that reduce the computational complexity, this Bayesian formulation allows for a more flexible separation of source influences onto the signal.

## Conclusion

In this research paper we extended a known framework to model motif influence on gene expression signal, which was originally introduced by The Fantom Consortium and the Riken Omics Science Center (2009); Balwierz *et al.* (2014). While the previous formulation assumes independence between samples, our Bayesian formulation provides the possibility to relax this assumption and allows to model correlation between samples. We first ran a simulation study on the basis of which we showed the superiority of the Bayesian Linear Mixed Model over Ridge Regression for data with independent noise. For noise that is dependent on the signal, the Bayesian Linear Mixed Model quickly loses its predictive power and has a similar performance to Ridge Regression. We further compared the two model assumptions on two real-world datasets: H3K27ac and RNA-seq data. We observed the same phenomenon as in the simulation study: no distinct superiority of the Bayesian Linear Mixed Model over Ridge Regression.

Despite the general theoretical improvement of the model, no clear dominance of that relaxation of the independence assumption between samples could be shown for data where the unexplained signal (noise) is highly correlated to the explained signal.

The advancements made in faster implementations together with mathematical reformulations, as done by Lippert *et al.* (2014a); Casale *et al.* (2015), allow for the usage of more complex models, such as the Bayesian Linear Mixed Model over simple Ridge Regression. In concert with the increase in computational power, such increase of mathematical complexity becomes more feasible to work with and no longer represents a practicality constraint as it used to.

Despite the drawback of the Bayesian Linear Mixed Model to not gain predictive power over Ridge Regression when applying it to data where large parts of the signal cannot be explained, we believe that further improvement of the formulation of the model’s covariates, e.g. motif counts from promoter and enhancer regions, will add value to the research community to better understand motif influences on gene expression.

Furthermore, we showed with the applications on the RNA-seq data that several signals of known motifs were picked up by Bayesian Linear Mixed Model whilst Ridge Regression ignored them and vice versa. This powerful model allows to take the interaction between samples into account to better understand the interplay between TFs and TF-mediated gene expression. It is yet a small step closer to the task of “decoding the genome the way TFs do” (Lambert *et al*., 2018).

## Acknowledgements

We thank Paolo Casale for his input and support on some technical details for the usage of the python package `limix` (Lippert *et al*., 2015).

## Authors’ Contribution

Concept: CA; Formal Analysis, Software and Writing: SL; Methodology: CA, SJvH, SL; Supervision: CA, SJvH, CA; Writing Review and Editing: SIL, SJvH, TH, CA.

## References

Anders, S. and Huber, W. (2010). Differential expression analysis for sequence count data. Genome Biol., 11(10), R106.

Balwierz, P. J. et al. (2014). ISMARA: automated modeling of genomic signals as a democracy of regulatory motifs. Genome Res., 24(5), 869–884.

Bollag, W. B. and Bollag, R. J. (2011). Wherefore art thou, YY1. J. Invest. Dermatol., 131(1), 11–12.

Bruse, N. and van Heeringen, S. J. (2018). GimmeMotifs : an analysis framework for transcription factor motif analysis. bioRxiv.

Casale, F. P. et al. (2015). Efficient set tests for the genetic analysis of correlated traits. Nat. Methods, 12(8), 755–758.

Collado-Torres, L. et al. (2017a). recount workflow: Accessing over 70,000 human RNA-seq samples with Bioconductor. F1000Research, 6(0), 1558.

Collado-Torres, L. et al. (2017b). Reproducible RNA-seq analysis using recount2. Nat. Biotechnol., 35, 319.

Creyghton, M. P. et al. (2010). Histone H3K27ac separates active from poised enhancers and predicts developmental state. Proc. Natl. Acad. Sci., 107(50), 21931–21936.

Ellis, S. E. et al. (2018). Improving the value of public RNA-seq expression data by phenotype prediction. Nucleic Acids Res., 46(9), e54–e54.

Elowitz, M. B. et al. (2002). Stochastic Gene Expression in a Single Cell. Science, 1183(297), 1183–1186.

Frankish, A. et al. (2019). GENCODE reference annotation for the human and mouse genomes. Nucleic Acids Res., 47(D1), D766–D773.

Geertz, M. et al. (2012). Massively parallel measurements of molecular interaction kinetics on a microfluidic platform. Proc. Natl. Acad. Sci., 109(41), 16540–16545.

Gunnell, A. et al. (2016). RUNX super-enhancer control through the Notch pathway by Epstein-Barr virus transcription factors regulates B cell growth. Nucleic Acids Res., 44(10), 4636–4650.

Hansen, K. D. et al. (2011). Sequencing technology does not eliminate biological variability. Nat. Biotechnol., 29(7), 572–573.

Huber, W. et al. (2003). Parameter estimation for the calibration and variance stabilization of microarray data. Stat. Appl. Genet. Mol. Biol., 2(1), Article 3.

Hunter, J. D. (2007). Matplotlib: A 2D Graphics Environment. Comput. Sci. Eng., 9(3), 90–95.

Irizarry, R. A. et al. (2005). Multiple-laboratory comparison of microarray platforms. Nat. Methods, 2(5), 345–350.

Koleti, A. et al. (2018). Data Portal for the Library of Integrated Network-based Cellular Signatures (LINCS) program: Integrated access to diverse large-scale cellular perturbation response data. Nucleic Acids Res., 46(D1), D558–D566.

Lambert, S. A. et al. (2018). The Human Transcription Factors. Cell, 172(4), 650–665.

Lambert, S. A. et al. (2019). Similarity regression predicts evolution of transcription factor sequence specificity. Nat. Genet., 51(6), 981–989.

Lee, T. I. and Young, R. A. (2013). Transcriptional Regulation and Its Misregulation in Disease. Cell, 152(6), 1237–1251.

Leek, J. T. et al. (2010). Tackling the widespread and critical impact of batch effects in high-throughput data. Nat. Rev. Genet., 11(10), 733–739.

Lieberman-Aiden, E. et al. (2009). Comprehensive Mapping of Long-Range Interactions Reveals Folding Principles of the Human Genome. Science, 326(5950), 289–293.

Lippert, C. et al. (2014a). LIMIX: genetic analysis of multiple traits. bioRxiv, pages 1–26.

Lippert, C. et al. (2014b). Supplemental Information Multivariate analysis of heritable traits. bioRxiv.

Lippert, C. et al. (2015). limix: linear mixed models for genomic analysis.

Lonsdale, J. et al. (2013a). The Genotype-Tissue Expression (GTEx) project. Nat. Genet., 45(6), 580–585.

Lonsdale, J. et al. (2013b). The Genotype-Tissue Expression (GTEx) project. Nat. Genet., 45(6), 580–585.

Madsen, J. G. S. et al. (2018). Integrated analysis of motif activity and gene expression changes of transcription factors. Genome Res., 28(2), 243–255.

Martens, J. H. A. and Stunnenberg, H. G. (2013). BLUEPRINT: mapping human blood cell epigenomes. Haematologica, 98(10), 1487–1489.

McKinney, W. (2010). Data Structures for Statistical Computing in Python. In S. van der Walt and J. Millman, editors, Proc. 9th Python Sci. Conf., pages 51–56.

Millman, K. J. and Aivazis, M. (2011). Python for Scientists and Engineers. Comput. Sci. Eng., 13(2), 9–12.

National Center for Biotechnology Information (US) (2019). Entrez-Gene: YY1 transcription factor [Homo sapiens (human)].

Ng, A. Y. (2004). Feature selection, L 1 vs. L 2 regularization, and rotational invariance. In Twenty-first Int. Conf. Mach. Learn. - ICML ‘04, page 78, New York, New York, USA. ACM Press.

Okuda, T. et al. (2001). RUNX1 / AML1 : A Central Player in Hematopoiesis. Int. J. Hematol., 74(3), 252–253.

Oliphant, T. E. (2006). Guide to Numpy. Trelgol Publishing USA, Austin, 2 edition.

Oliphant, T. E. (2007). Python for Scientific Computing. Comput. Sci. Eng., 9(3), 10–20.

Osmanbeyoglu, H. U. et al. (2014). Linking signaling pathways to transcriptional programs in breast cancer. Genome Res., 24(11), 1869–1880.

Pearson, K. (1895). VII. Note on regression and inheritance in the case of two parents. Proc. R. Soc. London, 58(347-352), 240–242.

Pedregosa, F. et al. (2011). Scikit-learn: Machine Learning in *{*P*}*ython. J. Mach. Learn. Res., 12, 2825–2830.

Qiao, M. et al. (2006). Cell Cycle-dependent Phosphorylation of the RUNX2 Transcription Factor by cdc2 Regulates Endothelial Cell Proliferation. J. Biol. Chem., 281(11), 7118–7128.

Quinlan, A. R. and Hall, I. M. (2010). BEDTools: a flexible suite of utilities for comparing genomic features. Bioinformatics, 26(6), 841–842.

Rakitsch, B. et al. (2013). It is all in the noise: Efficient multi-task Gaussian process inference with structured residuals. In C. Burges, L. Bottou, M. Welling, Z. Ghahramani, and K. Weinberger, editors, Adv. Neural Inf. Process. Syst. 26 (NIPS 2013), pages 1466–1474.

Schmidt, F. et al. (2017). Combining transcription factor binding affinities with openchromatin data for accurate gene expression prediction. Nucleic Acids Res., 45(1), 54–66.

Shi, L. et al. (2006). The MicroArray Quality Control (MAQC) project shows inter- and intraplatform reproducibility of gene expression measurements. Nat. Biotechnol., 24(9), 1151–1161.

Sood, R. et al. (2017). Role of RUNX1 in hematological malignancies. Blood, 129(15), 2070–2082.

Spender, L. C. et al. (2002). Expression of Transcription Factor AML-2 (RUNX3, CBF -3) Is Induced by Epstein-Barr Virus EBNA-2 and Correlates with the B-Cell Activation Phenotype. J. Virol., 76(10), 4919–4927.

Stegle, O. and Lawrence, N. D. (2012). Joint Modelling of Confounding Factors and Prominent Genetic Regulators Provides Increased Accuracy in Genetical Genomics Studies. PLOS Comput. Biol., 8(1), 1–8.

Taguchi, S. et al. (2011). Overexpression of the transcription factor yin-yang-1 suppresses differentiation of HaCaT cells in three-dimensional cell culture. J. Invest. Dermatol., 131, 37–45.

Takahashi, K. and Yamanaka, S. (2006). Induction of Pluripotent Stem Cells from Mouse Embryonic and Adult Fibroblast Cultures by Defined Factors. Cell, 126(4), 663–676.

The Fantom Consortium and the Riken Omics Science Center (2009). The transcriptional network that controls growth arrest and differentiation in a human myeloid leukemia cell line. Nat. Genet., 41(5), 553–562.

Tibshirani, R. (1988). Estimating Transformations for Regression via Additivity and Variance Stabilization. J. Am. Stat. Assoc., 83(402), 394–405.

Tikhanovich, I. et al. (2013). FOXO Transcription Factors in Liver Function and Disease. J Gastroenterol Hepatol, 28(1), 125–131.

Uhlen, M. et al. (2017). A pathology atlas of the human cancer transcriptome. Science, 357(660), 1–11.

van Heeringen, S. J. and Veenstra, G. J. C. (2011). GimmeMotifs: a de novo motif prediction pipeline for ChIP-sequencing experiments. Bioinformatics, 27(2), 270–271.

Waskom, M. et al. (2017). mwaskom/seaborn: v0.8.1 (September 2017).

Wickham, H. (2016). ggplot2. Cham.

Wilke, C. O. (2018). cowplot: Streamlined Plot Theme and Plot Annotations for ‘ggplot2’.

