## Supplementary material for "Bayesian Linear Mixed Models for Motif Activity Analysis": Supplelemt Material

September 25, 2019

#### A Bayesian Linear Mixed Models

In this section, we explain in more detail the Bayesian take on the linear model

$$\bar{\mathbf{y}}_{g,c} = \sum_{t=1}^{T} \mathbf{m}_{t,g}\omega_{t,c} + \text{noise}, \tag{1}$$

where we model the normalized gene expression signal $\bar{\mathbf{y}}_{g,c} = \mathbf{y}_{g,c} - \bar{\mathbf{y}}_g - \bar{\mathbf{y}}_c$ as a linear combination of the motif scores $\mathbf{M}_{T,G}$, which are each, dependent on the sample $c$ and the TF $t$, weighted with $\omega_{T,C}$. The term 'noise' represents all signal that cannot be explained by the model, i.e. the linear combination of the motif scores $\mathbf{M}_{T,G}$. This can be any technical noise, motif influence for which the linear assumption might be too simplistic, but also any other source that drives the gene expression $\mathbf{y}_{g,c}$ and is not modeled. The model was originally introduced by The Fantom Consortium and the Riken Omics Science Center (2009) and subsequently expanded by Balwierz *et al.* (2014).

The main idea behind a Bayesian approach is to include some prior knowledge into the data, called the prior. Here, we model $\omega_{t,c}$, the influence of motif $t \in \{1,\ldots,T\}$ in condition $c \in \{1,\ldots,C\}$, as a normal distributed prior with mean zero. Its marginal distributions are

$$\omega_C \sim \mathcal{N}(\mathbf{0}, \mathbf{V}_C), \tag{2}$$

where $\mathbf{V}_C$ is the covariance in motif activity over all conditions and

$$\omega_T \sim \mathcal{N}\left(\mathbf{0}, \sigma^{\mathbf{2}}\mathbf{I}_T\right), \tag{3}$$

with $\sigma^{\mathbf{2}}\mathbf{I}_T$ being the covariance over all motifs. Hence, we assume independence between motifs. Assuming dependence between motifs with covariance $\mathbf{\Psi}$ can easily be implemented in the model. Making use of the vectorization representation of multivariate normal distribution, we can write the multivariate Normal distribution of $\omega_{T,C}$ as follows:

$$\text{vec}(\omega_{T,C}) \sim \mathcal{N}\left(\mathbf{0}, \sigma^{\mathbf{2}}\mathbf{V}_C \otimes \mathbf{I}_T\right). \tag{4}$$

Analogously, we can rewrite Eq. 1 in matrix-vector notation:

$$\text{vec}(\mathbf{Y}_{G,C}) \sim \mathcal{N}\left(\mathbf{M}_{T,G}^{\intercal}\omega_{T,C}, \delta\mathbf{\Sigma}_C \otimes \mathbf{I}_G\right), \tag{5}$$

with $\mathbf{\Sigma}_C$ being the covariance conditions and $\delta\mathbf{I}_G$ covariance over genes. In the following, we use Bayes' rule on marginal and conditional Gaussians (see Bischop (2006, Chapter 2.3, p. 93)). We first list the general formulas here, which we copy from Bischop (2006): The marginal distribution of $\mathbf{x}$ and conditional distribution of $\mathbf{y}$ given $\mathbf{x}$ are given as follows:

$$p(\mathbf{x}) = \mathcal{N}\left(\mathbf{x}|\mu, \mathbf{\Lambda}^{-1}\right) \tag{6}$$

$$p(\mathbf{y}|\mathbf{x}) = \mathcal{N}\left(\mathbf{y}|\mathbf{A}\mathbf{x} + \mathbf{b}, \mathbf{L}^{-1}\right). \tag{7}$$

Translating that notation into our own notation, we get the following equivalences:

$$\mathbf{x} \equiv \mathrm{vec}\left(\omega_{T,C}\right) \qquad \mu \equiv \mathbf{0} \qquad \mathbf{\Lambda}^{-1} \equiv \sigma^{\mathbf{2}}\mathbf{V}_C \otimes \mathbf{I}_T \tag{8}$$

$$\mathbf{y} \equiv \mathrm{vec}\left(\mathbf{Y}_{G,C}\right) \qquad \mathbf{A} \equiv \mathbf{M}_{T,G}^{\mathbf{T}} \qquad \mathbf{b} \equiv \mathbf{0} \qquad \mathbf{L}^{-1} \equiv \delta\mathbf{\Sigma}_C \otimes \mathbf{I}_G. \tag{9}$$

In our own notation, and following Eq. 7, the distribution of $\mathbf{Y}_{G,C}$ given $\omega_{T,C}$ can be rewritten as:

$$\mathrm{vec}\left(\mathbf{Y}_{G,C}|\omega_{T,C}\right) \sim \mathcal{N}\left(\mathbf{0}, \sigma^{\mathbf{2}}\mathbf{V}_C \otimes \mathbf{\Pi}_G + \delta\mathbf{\Sigma}_C \otimes \mathbf{I}_G\right), \tag{10}$$

where $\mathbf{\Pi}_G = \mathbf{M}_{T,G}^{\mathbf{T}}\mathbf{M}_{T,G}$.

Going back to the general formula from Bischop (2006), the marginal distribution of $\mathbf{y}$ and conditional distribution of $\mathbf{x}$ given $\mathbf{y}$ are computed according to the following:

$$p(\mathbf{y}) = \mathcal{N}\left(\mathbf{y}|\mathbf{A}\mu + \mathbf{b}, \mathbf{L}^{-1} + \mathbf{A}\mathbf{\Lambda}^{-1}\mathbf{A}^T\right) \tag{11}$$

$$p(\mathbf{x}|\mathbf{y}) = \mathcal{N}\left(\mathbf{x}|\mathbf{\Sigma}\left\{\mathbf{A}^T\mathbf{L}\left(\mathbf{y} - \mathbf{b}\right) + \mathbf{\Lambda}\mu\right\}, \mathbf{\Sigma}\right), \tag{12}$$

$$\text{with } \mathbf{\Sigma} = \left(\mathbf{\Lambda} + \mathbf{A}^T\mathbf{\Lambda}\mathbf{A}\right)^{-1}. \tag{13}$$

Hence, based on Eq. 11 and Eq. 12, the posterior distribution of $\omega_{T,C}$ given $\mathbf{Y}_{G,C}$ is then

$$\mathrm{vec}\left(\omega_{T,C}|\mathbf{Y}_{G,C}\right) \sim \mathcal{N}\left(\left(\mathbf{V}_C \otimes \mathbf{M}_{T,G}^{\mathbf{T}}\mathbf{I}_T\right)\mathbf{\Lambda}_{\mathbf{CG}\otimes\mathbf{CG}}{}^{-1}\mathrm{vec}\left(\mathbf{Y}_{G,C}\right), \mathbf{\Lambda}_{\mathbf{CG}\otimes\mathbf{CG}}\right) \tag{14}$$

with

$$\mathbf{\Lambda}_{\mathbf{CG}\otimes\mathbf{CG}} = \sigma^{\mathbf{2}}\mathbf{V}_C \otimes \mathbf{\Pi}_G + \delta\mathbf{\Sigma}_C \otimes \mathbf{I}_G. \tag{15}$$

For the reformulation, we used the *Woodbury matrix identity* (Woodbury, 1950). For the computation of the posterior values of $\omega_{T,C}$, with notation $\hat{\omega}_{T,C}$, it suffices to compute the mean of Eq. 14:

$$\mathrm{vec}\left(\hat{\omega}_{T,C}|\mathbf{Y}_{G,C}\right) = \left(\mathbf{V}_C \otimes \mathbf{M}_{T,G}^{\mathbf{T}}\mathbf{I}_T\right)\left[\sigma^{\mathbf{2}}\mathbf{V}_C \otimes \mathbf{\Pi}_G + \delta\mathbf{\Sigma}_C \otimes \mathbf{I}_G\right]^{-1}\mathrm{vec}\left(\mathbf{Y}_{G,C}\right) \tag{16}$$

As the covariance matrix is assumed to be the sum of Kronecker products, the runtime complexity is reduced to $O\left(G^3 + C^3\right)$ in a $O\left(G^2 + C^2\right)$ space instead of $O\left(G^3C^3\right)$ runtime and a memory requirement of $O\left(G^2C^2\right)$ (Lippert *et al.*, 2014).

### B Simulating data

For the simulation study we generate data based on the model introduced in Eq. 1-Eq. 4, given a covariance matrix $\mathbf{V}_C$. The prior weight $\tilde{\omega}_{T,C}$ and $\mathbf{Y}_{G,C}$ are generated according to Eq. 2 and Eq. 1. The expression data $\mathbf{Y}_{G,C}$ from Eq. 1 is then used to estimate $\mathbf{V}_C$ and $\mathbf{\Sigma}_C$. Based on these computations, the posterior motif influence $\hat{\omega}_{T,C}$ is then computed and compared to the simulated motif influence $\tilde{\omega}_{T,C}$ with a Pearson correlation (Pearson, 1895) over all conditions. As gene set we use the 978 landmark genes from the LINCS project (Koleti *et al.*, 2018). In a secondary simulation we increase the size of the gene set to 5000 genes, which originate from an analysis of the most variational genes across all samples from the GTEx project (Genotype Tissue Expression, `https://gtexportal.org/home/`). We generate data for $C = \{10, 30, 50, 70, 100, 120\}$ conditions.

#### Covariance types of $\mathbf{V}_C$

For the generation of simulated motif influence $\tilde{\omega}_{T,C}$ (Eq. 2), a covariance matrix $\sigma^{\mathbf{2}}\mathbf{V}_C \otimes \mathbf{I}_T$ needs to be given. As the covariance along TFs is modeled to be independent, one can generate $\omega_{t,C}$ randomly $T$ times with $\omega_{t,C} \sim \mathcal{N}\left(0, \sigma^{\mathbf{2}}\mathbf{V}_C\right)$. In the following, four covariance types are explained that we generate as $\mathbf{V}_C$ to serve as covariance for the generated $\omega_{T,C}$ in Eq. 2 and hence expression data $\mathbf{Y}_{G,C}$ in Eq. 3. All covariance matrices are then normalized by their trace. Below, we explain how $\sigma^{\mathbf{2}}$ is generated.

##### Independence

For independent conditions, we simply set $\mathbf{V}_C$ to an identity matrix: $\mathbf{V}_C = \sigma^{\mathbf{2}}\mathbf{I}_C$. We refer to these as independent, but in reality they are even isotropic, as the covariance between samples is assumed to be identical.

##### Unrestricted correlation

A random (uniform) $C \times C$ matrix $\mathbf{R}_C$ is generated and added with an identity matrix. This is then multiplied with itself: $\mathbf{V}_C = \sigma^{\mathbf{2}}\left(\mathbf{I}_C + \mathbf{R}_C\right)\left(\mathbf{I}_C + \mathbf{R}_C\right)^{\mathsf{T}}$

##### Correlation - high correlation and 50% correlation

We assume correlation between samples to allow for replicates and similarities between cell-lines. Hence, we would expect that those samples cluster in blocks when applying a clustering algorithm to the data (see Fig. 6 or Fig. 11 as examples). Therefore, we generate these block matrices with $k$ blocks of ones. Samples within are block are then completely correlated and completely independent from samples outside that block. The size of the blocks is minimally one and randomly determined. The sum of all block sizes is $C$. These blocks are lined up along the diagonal. The non-block elements are set to $-0.01$. The procedure is written in pseudo-code in Supplement B. For an example of a $k = \frac{1}{2}C$ covariance matrix, see Fig. 2A and B, left panel. For the analysis, we generate highly correlated covariance matrices with two blocks, i.e. $k = 2$, and 50% correlated covariance matrices, they are hence of medium rank, with $k = \frac{1}{2}C$ correlated sample groups.

#### Noise $\mathbf{\Sigma}_C$ - unstructured and structured

To generate the gene expression data $\mathbf{Y}_{G,C}|\omega_{T,C}$, we compute the signal as the product of the motif scores $\mathbf{M}_{T,G}$ (explained hereafter) and $\omega_{T,C}$ (explained previously). Due to the randomness in the signal that is not explained by motifs, we add some random noise which is drawn from a normal distribution with covariance $\delta\mathbf{\Sigma}_C \otimes \mathbf{I}_G$. We generate $\mathbf{\Sigma}_C$ in two different ways: (i) assuming no particular structure, $\mathbf{\Sigma}_{C,\text{random}}$, which is a matrix filled with values drawn from a standard normal distribution, multiplied with itself, or (ii) adding structure that is similar to $\mathbf{V}_C$: $\mathbf{\Sigma}_{C,\mathbf{V}_C,\rho} = \mathbf{\Sigma}_{C,\text{random}} + \eta_\rho \mathbf{V}_C$. The noise matrices are then normalized by their trace.

##### Unstructured noise $\mathbf{\Sigma}_C$

In a first step, we randomly generate a covariance matrix, analogously to the random covariance matrix $\mathbf{V}_C$: by taking the product of a randomly drawn matrix, we assure its symmetry and positive definiteness. A visualization is given in Fig. 2A, right panel.

##### Structured noise $\mathbf{\Sigma}_C$

To the unstructured noise matrix $\mathbf{\Sigma}_C$, explained in the previous paragraph, we add a structure that is similar to $\mathbf{V}_C$:

$$\mathbf{\Sigma}_{C,\mathbf{V}_C,\rho} = \eta_\rho \mathbf{V}_C + \mathbf{\Sigma}_{C,\text{random}}, \tag{17}$$

where $\eta_\rho$ depends on $\rho \in [0, 1]$, the degree of structure in the noise. If $\rho = 0$, the noise is unstructured, and a fraction $\rho$ of one yields the sum of $\mathbf{V}_C$ and $\mathbf{\Sigma}_C$, normalized by their traces, respectively:

$$\eta_\rho = \begin{cases} 0 & \text{if } \rho = 0 \\ \frac{\rho}{1-\rho} \frac{\text{tr}(\mathbf{\Sigma}_C)}{\text{tr}(\mathbf{V}_C)} & \text{for } \rho \in (0, 1) \end{cases} \tag{18}$$

$$\mathbf{\Sigma}_{C,\mathbf{V}_C,\rho} = \eta_\rho \mathbf{V}_C + \mathbf{\Sigma}_{C,\text{random}} \tag{19}$$

We therefore control the degree of structure in the noise. A visualization of such a structured noise matrix is given in Fig. 2B, right panel. As any model is just a simplification of reality, we cannot explain the entire signal expressed in the expression data $\mathbf{Y}_{G,C}$. This is especially the case as we model the signal uniquely as a linear product of artificially computed motif scores. Hence, there will always be signal in the data that cannot be explained by motifs. We therefore add to the random environmental or technical noise signal that explains the relationship between conditions.

##### Signal-to-noise ratio

From previous research (Balwierz *et al.*, 2014), it has been shown that roughly $10 - 20\%$ of the signal of gene expression can be explained by motif influence in the promoter region. We therefore generate the data in such a way, that 20% of the signal in expression data $\mathbf{Y}_{G,C}$ is due to motifs, and the rest unexplainable noise. We achieve this by adjusting the parameter $\boldsymbol{\sigma^2}$ and $\delta$. The latter is fixed by the rough percentage wished to be expressed by the noise, $1 - \beta$, which we set to $\beta = 0.2$. For scaling reasons, it is divided with the 2-norm of $\mathbf{\Sigma}_C$:

$$\delta = \frac{(1 - \beta)}{\|\mathbf{\Sigma}_C\|_2}, \tag{20}$$

with $\|\cdot\|_2 = \sigma_{\max}(\cdot)$ as the matrix 2-norm which is equivalent to $\sigma_{\max}(\cdot)$, the maximal singular value. $\boldsymbol{\sigma^2}$ is determined by bisection, such that it explains 0.2 of the variance coefficient (Eq. 1). As starting value, we set:

$$\boldsymbol{\sigma^2} = \delta \frac{\beta}{1 - \beta} \frac{C \text{tr}(\mathbf{\Sigma}_C)}{\text{tr}(\mathbf{V}_C) \, \text{tr}(\mathbf{\Pi}_G)}. \tag{21}$$

#### Pseudocode for the generation of lower rank block matrices

**Algorithm 1** Generation of correlated covariance matrices

```
function generateBlocksizes( C, k )          ▷ generate k blocksizes that sum up to C
    num_blocks ← k
    blocksizes ← initialize vector of length num_blocks
    for block_i in (num_blocks − 1) do
        lenght_blocks ← sum( blocksizes )
        leftover-space ← C − length_blocks + 1
        blocksize_i ← random integer between 1 and leftover-space
        blocksizes[i] ← blocksize_i
    blocksizes[num_blocks] ← C − block-lengths + 1
    return blocksizes
procedure Block matrix of dimension C with k blocks of size blocksizes
    matrix ← initialize matrix of dimension C
    block_i_start ← 1
    block_i_end ← 0
    blocksizes ← generateBlocksizes(C, k)
    for blocksize_i in blocksizes do
        block_i_end ← block_i_end + blocksize_i
        matrix[block_i_start : block_i_end, block_i_start : block_i_end] ←
            matrix of ones of size blocksize_i      ▷ place matrix of ones onto diagonal
        block_i_start ← block_i_end
    for all elements in matrix that are zero do
        fill with 1e−2                              ▷ fill all off-diagonal elements
```

### References


Balwierz, P. J. *et al.* (2014). ISMARA: automated modeling of genomic signals as a democracy of regulatory motifs. *Genome Res.*, **24**(5), 869–884.

Bischop, C. M. (2006). *Pattern Recognition and Machine Learning*. Springer Science+Business Media, 9 edition.

Koleti, A. *et al.* (2018). Data Portal for the Library of Integrated Network-based Cellular Signatures (LINCS) program: Integrated access to diverse large-scale cellular perturbation response data. *Nucleic Acids Res.*, **46**(D1), D558–D566.

Lippert, C. *et al.* (2014). Supplemental Information Multivariate analysis of heritable traits. *bioRxiv*.

Pearson, K. (1895). VII. Note on regression and inheritance in the case of two parents. *Proc. R. Soc. London*, **58**(347-352), 240–242.

The Fantom Consortium and the Riken Omics Science Center (2009). The transcriptional network that controls growth arrest and differentiation in a human myeloid leukemia cell line. *Nat. Genet.*, **41**(5), 553–562.

Woodbury, M. A. (1950). Inverting modified matrices. *Memo. Rep.*, **42**(106), 336.

#### C Supplemental Figures

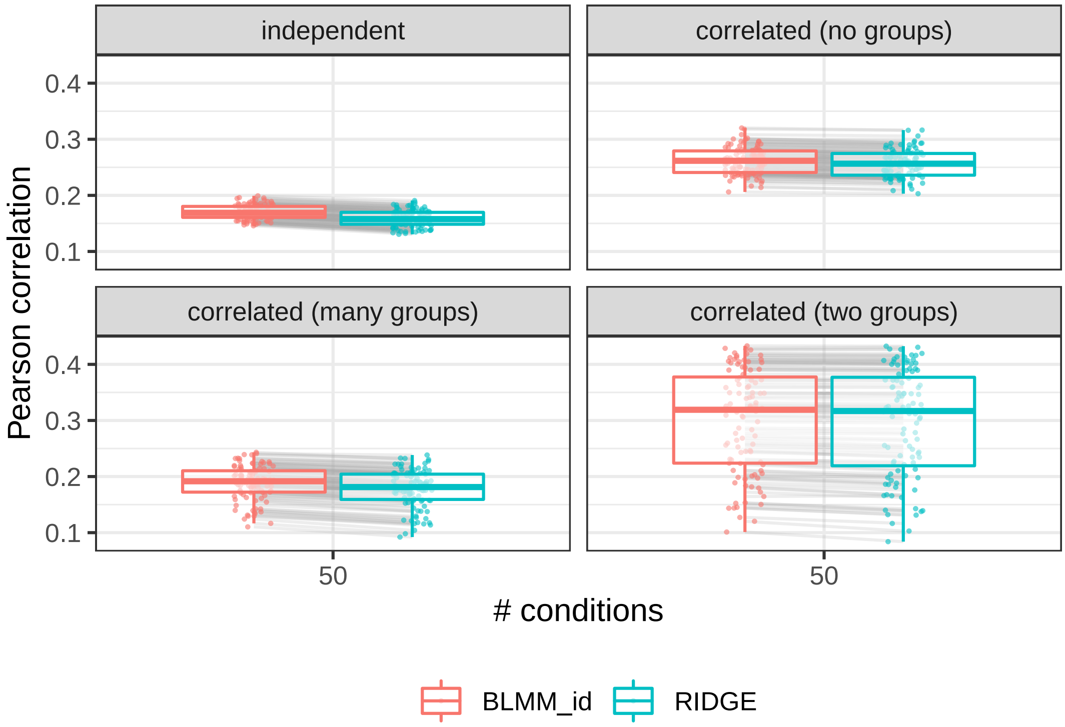


Figure 1: **Ridge Regression is a special case of Bayesian Linear Mixed Model, limiting the estimated covariance and noise to be independent.** Data was generated over $G = 978$ informative genes and $T = 623$ motif scores and 100 repetitions, with structured $\mathbf{\Sigma}_C$. The results for Bayesian Linear Mixed Model and Ridge Regression (RIDGE, in blue) are equal when Bayesian Linear Mixed Model is limited to $\mathbf{V}_C = \sigma^2\mathbf{I}_C$ and $\mathbf{\Sigma}_C = \delta\mathbf{I}_C$ (BLMM_id, in red). The Pearson correlation values are computed between the generated and predicted posterior motif influence $\hat{\omega}_{T,C}$, and separated by method used to compute them. Data is generated with (i) independent samples, $\mathbf{V}_C = \mathbf{I}_C$, (ii) unrestricted correlation between samples, (iii) 50% correlated data, where the samples cluster in many ($k = \frac{1}{2}C$) sample groups, (iv) highly correlated data, by generating a covariance matrix with $k = 2$ completely correlated sample groups, with $C$ the number of conditions.

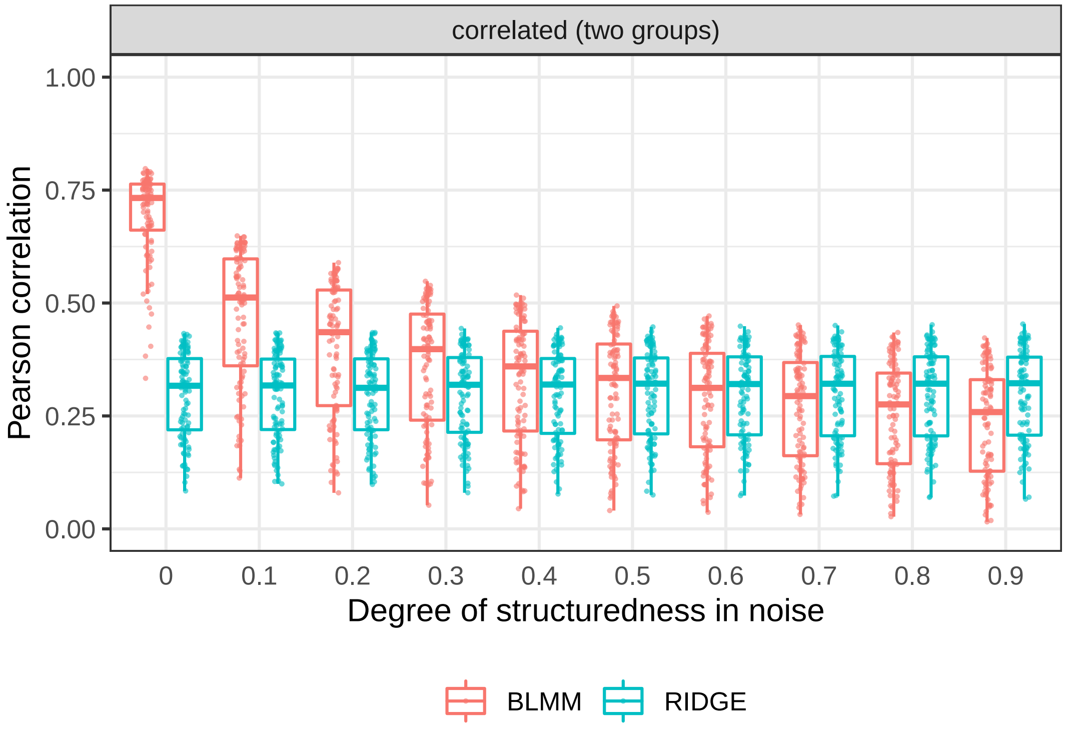


Figure 2: **Models' performance interchange superiority for data generated with increasing structuredness in noise** Data was generated over $G = 978$ informative genes, $T = 623$ motif scores, 100 repetitions, and with a covariance matrix "$\mathbf{V}_C$: highly correlated (two groups)". $\mathbf{\Sigma}_C$ is generated with increasing degree of structuredness (x-axis). Model performance of Bayesian Linear Mixed Model (BLMM, in red) and Ridge Regression (RIDGE, in blue) are shown on the bases of the Pearson correlation values between generated and predicted motif-condition-weights.

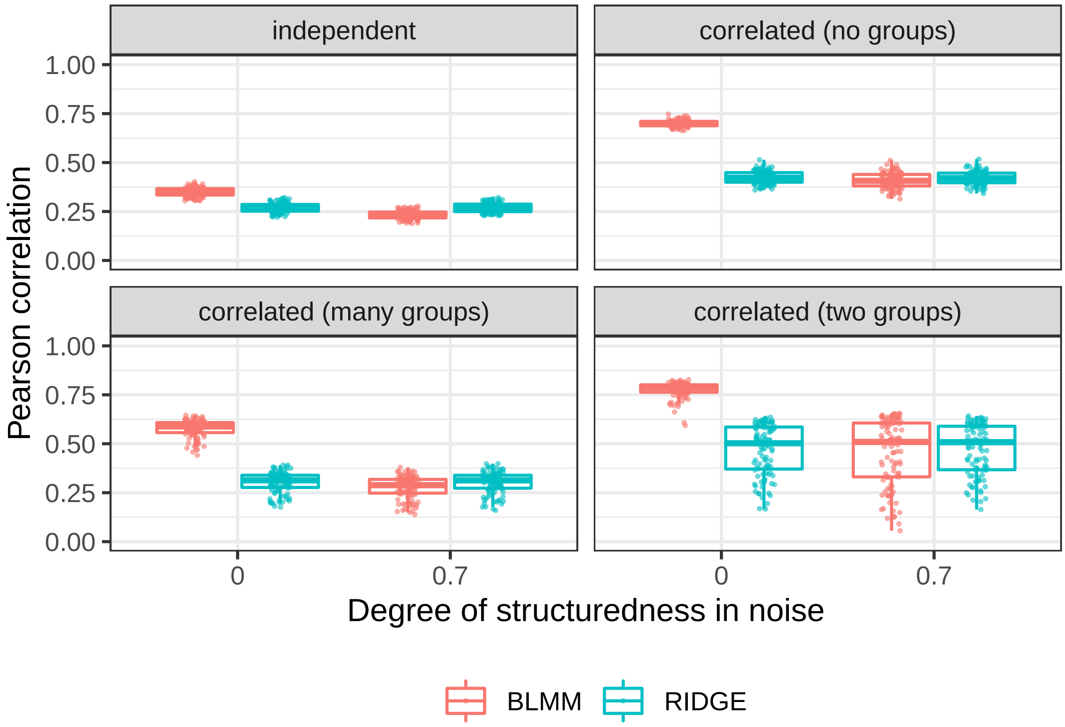


Figure 3: **Simulation study on $G = 5000$ genes** We compare the model assumptions of Bayesian Linear Mixed Model (BLMM, depicted in red) and Ridge Regression (LIMIX, in blue) on datasets generated with $C = 50$ samples, and $G = 5000$ genes. On the x-axis we show data that is generated with unstructured noise (degree of structuredness=0) and with structured noise with $\rho = 0.7$. On the y-axis, we depict the Pearson correlation values between generated and predicted motif-condition weights. In each panel, the data was generated with different assumptions on the degree of correlation between samples: (i) independence ($\mathbf{V}_C = \mathbf{I}_C$ (upper left), (ii) unrestricted correlation (upper right), (iii) correlated with many sample groupgs (lower left), and (iv) highly correlated with two sample groups) (lower right). There is no difference in performance when increasing the dimensionality of genes. The Bayesian Linear Mixed Model has predictive power over Ridge Regression when the data is correlated, uniquely for unstructured noise. For structured noise ($\rho = 0.7$), there is no gain in performance, despite the bigger size of the dataset.

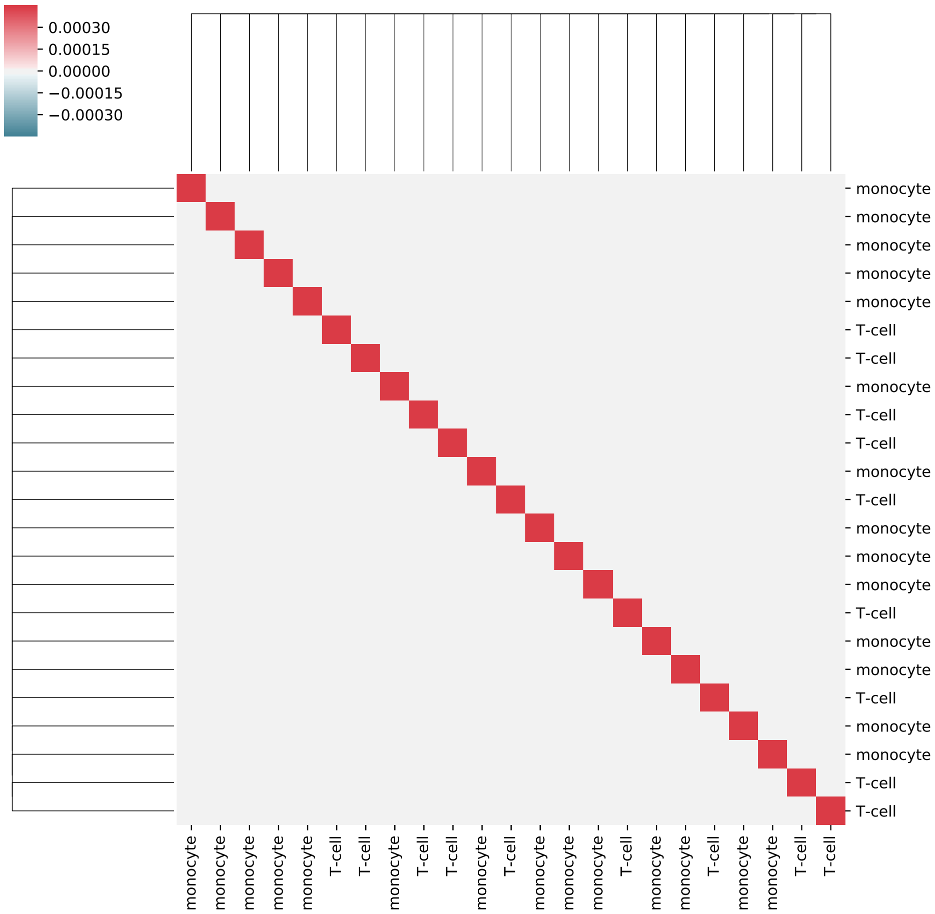


Figure 4: **H3K27ac: $\mathbf{V}_C$ for Ridge Regression** Estimated correlation between conditions $\mathbf{V}_C$ assuming independence between the conditions for the H3K27ac dataset.

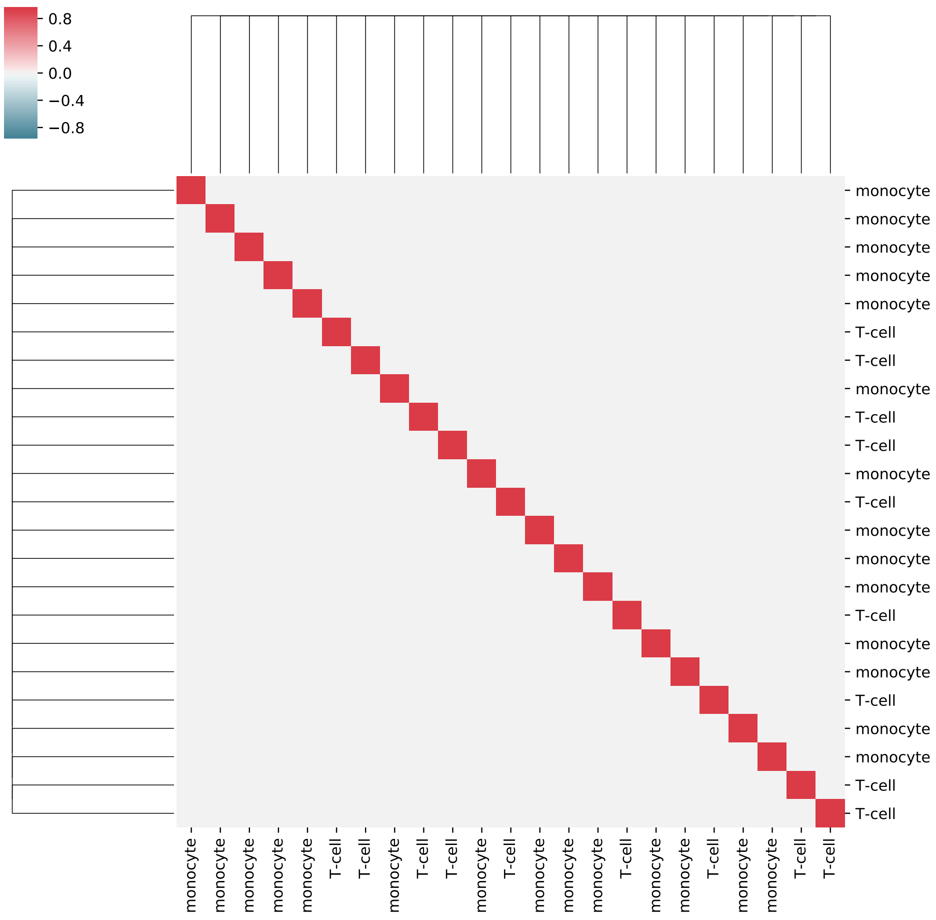


Figure 5: **H3K27ac: $\boldsymbol{\Sigma}_C$ for Ridge Regression** Estimated noise $\boldsymbol{\Sigma}_C$ assuming independence between the conditions for the H3K27ac dataset.

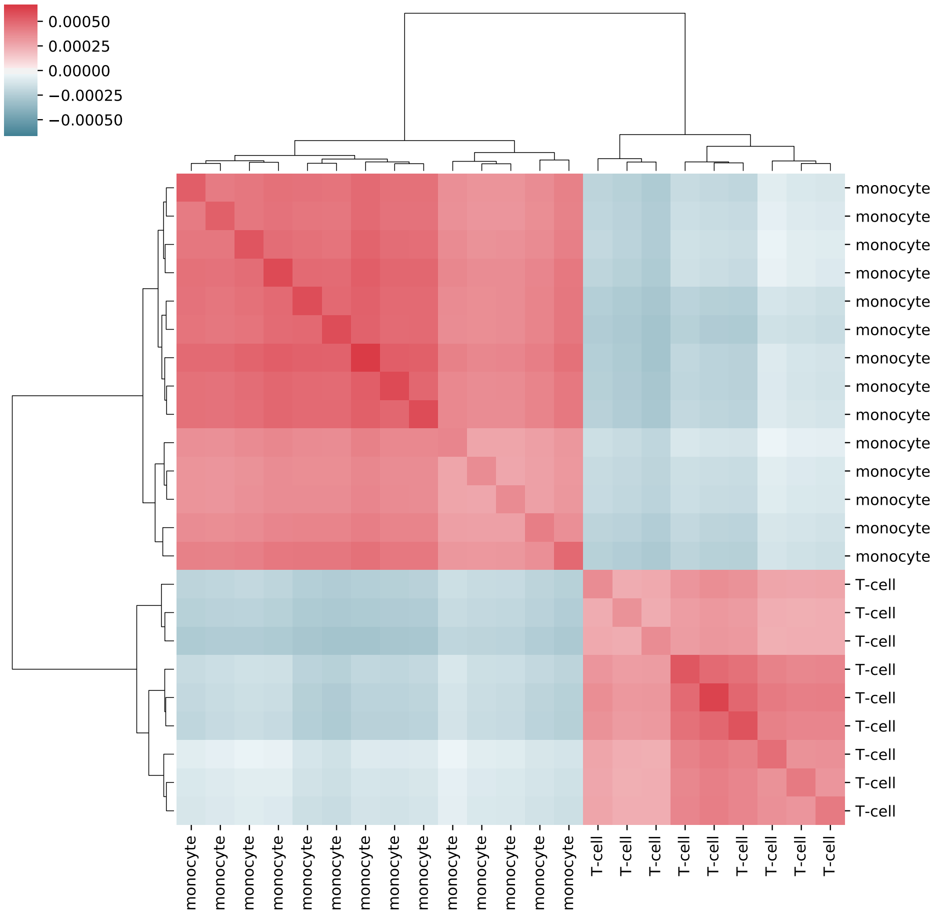


Figure 6: **H3K27ac: $\mathbf{V}_C$ for Bayesian Linear Mixed Model** Estimated correlation between conditions $\mathbf{V}_C$ assuming dependence between the conditions for the H3K27ac dataset.

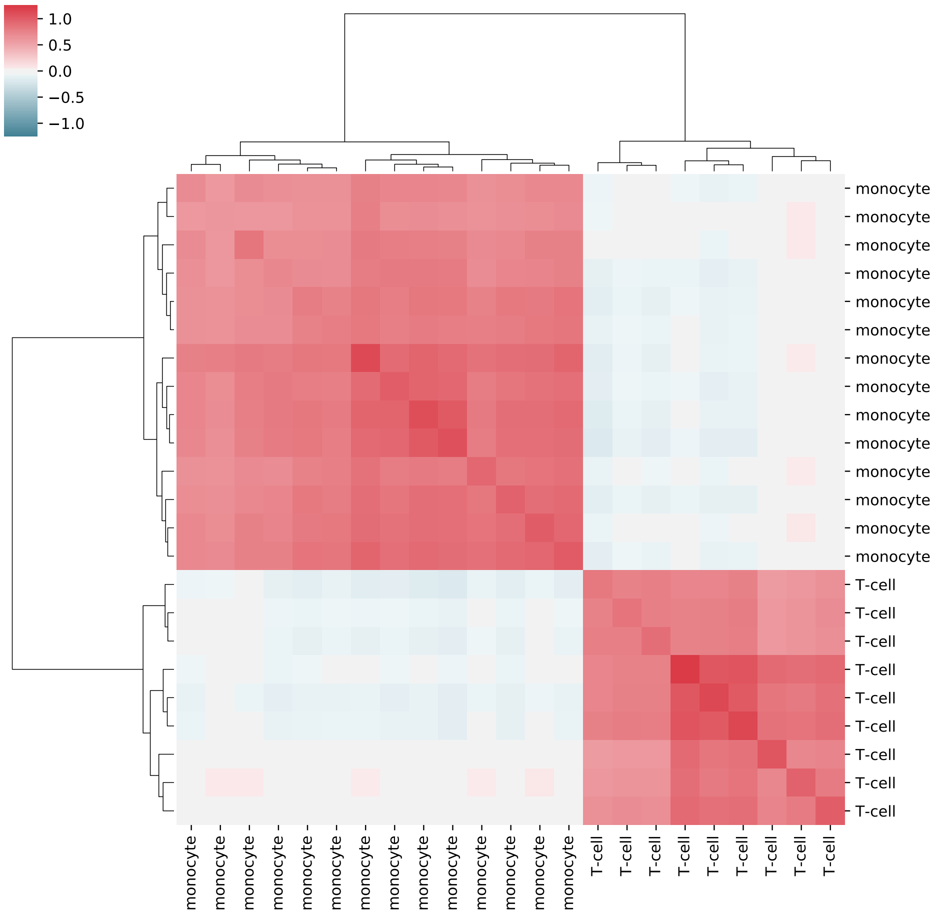


Figure 7: **H3K27ac: $\mathbf{\Sigma}_C$ for Bayesian Linear Mixed Model** Estimated noise $\mathbf{\Sigma}_C$ assuming dependence between the conditions for the H3K27ac dataset.

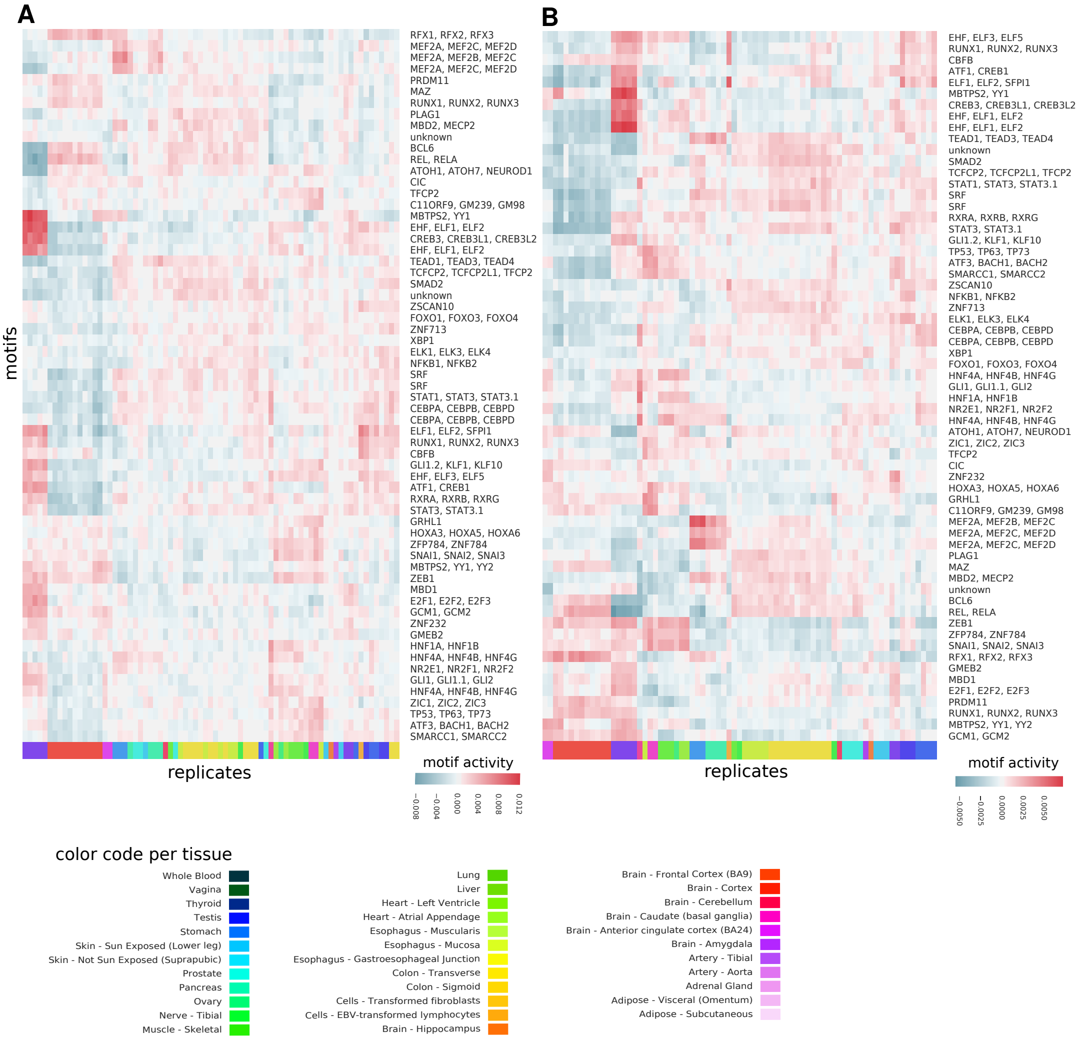


Figure 8: **Heatmaps GTEx motif-condition-weight matrix** Weights for the motif-condition-weights of those 56 common motifs, that are outside of 2.5 times the inter-quantile range for all motifs over a tissue. The weights are computed assuming dependence (A) or independence (B) between the conditions.

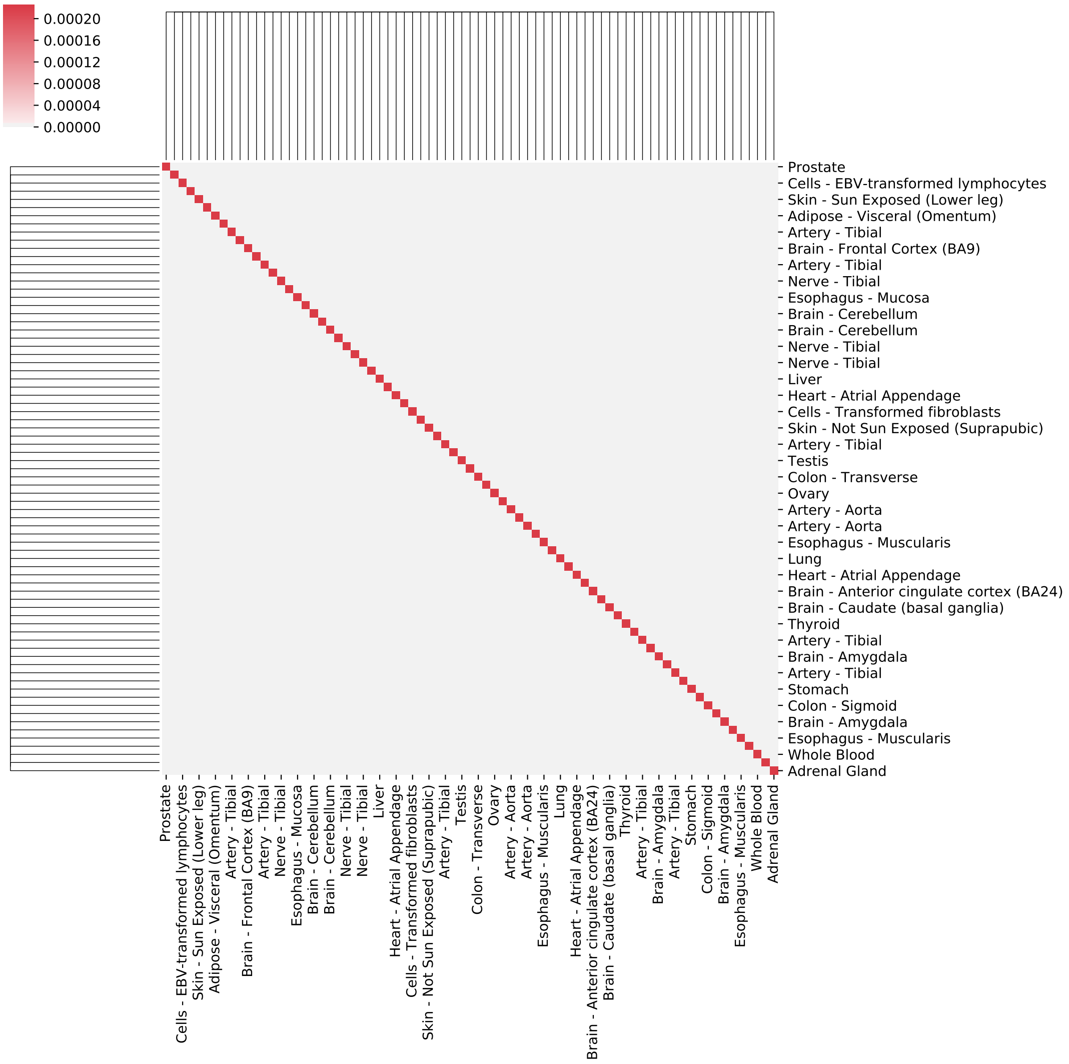


Figure 9: **GTEx: $\mathbf{V}_C$ for Ridge Regression** Estimated correlation between conditions $\mathbf{V}_C$ assuming independence between the conditions for the GTEx dataset.

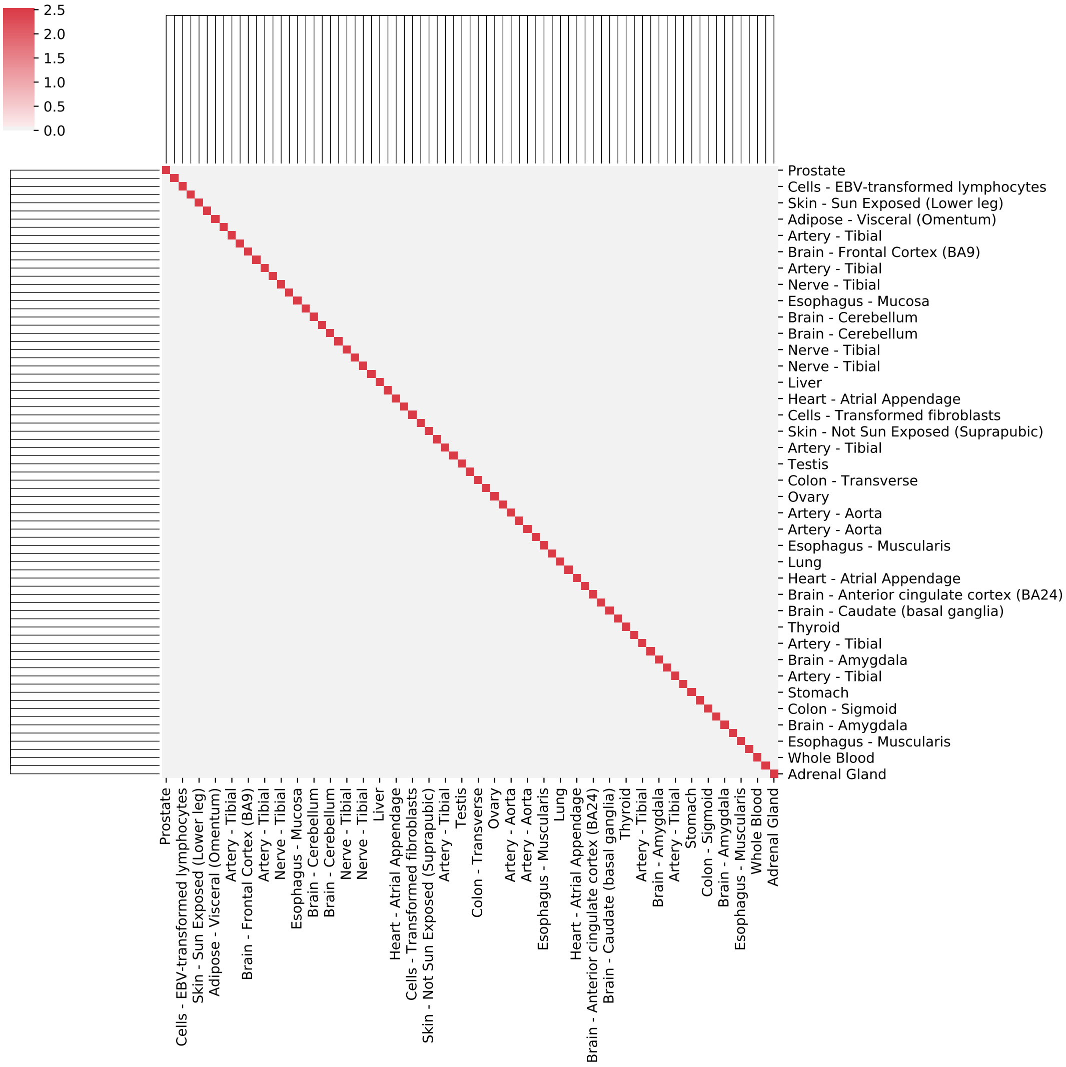


Figure 10: **H3K27ac: $\mathbf{\Sigma}_C$ for Ridge Regression** Estimated noise $\mathbf{\Sigma}_C$ assuming independence between the conditions for the GTEx dataset.

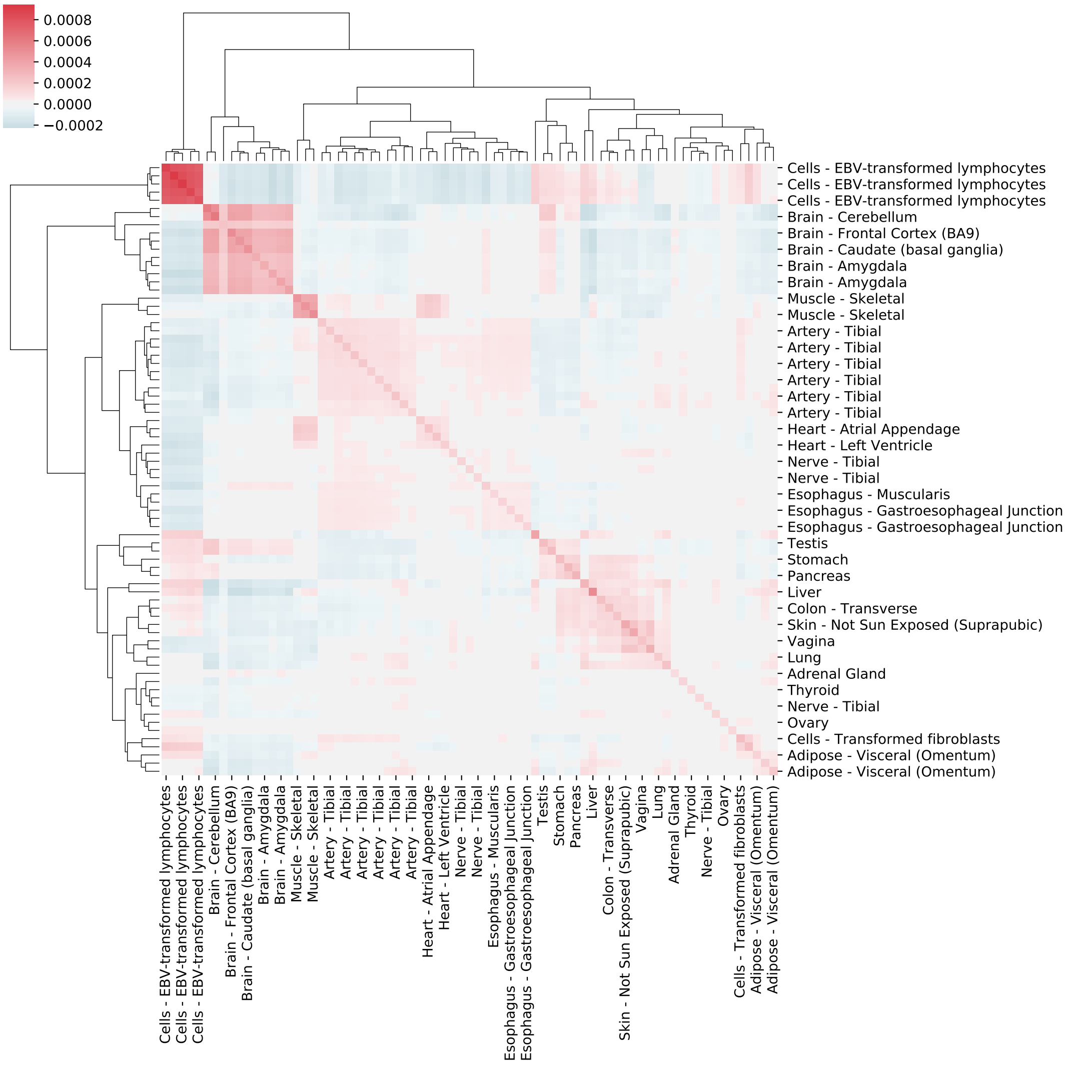


Figure 11: **GTEx: $\mathbf{V}_C$ for Bayesian Linear Mixed Model** Estimated correlation between conditions $\mathbf{V}_C$ assuming dependence between the conditions for the GTEx dataset.

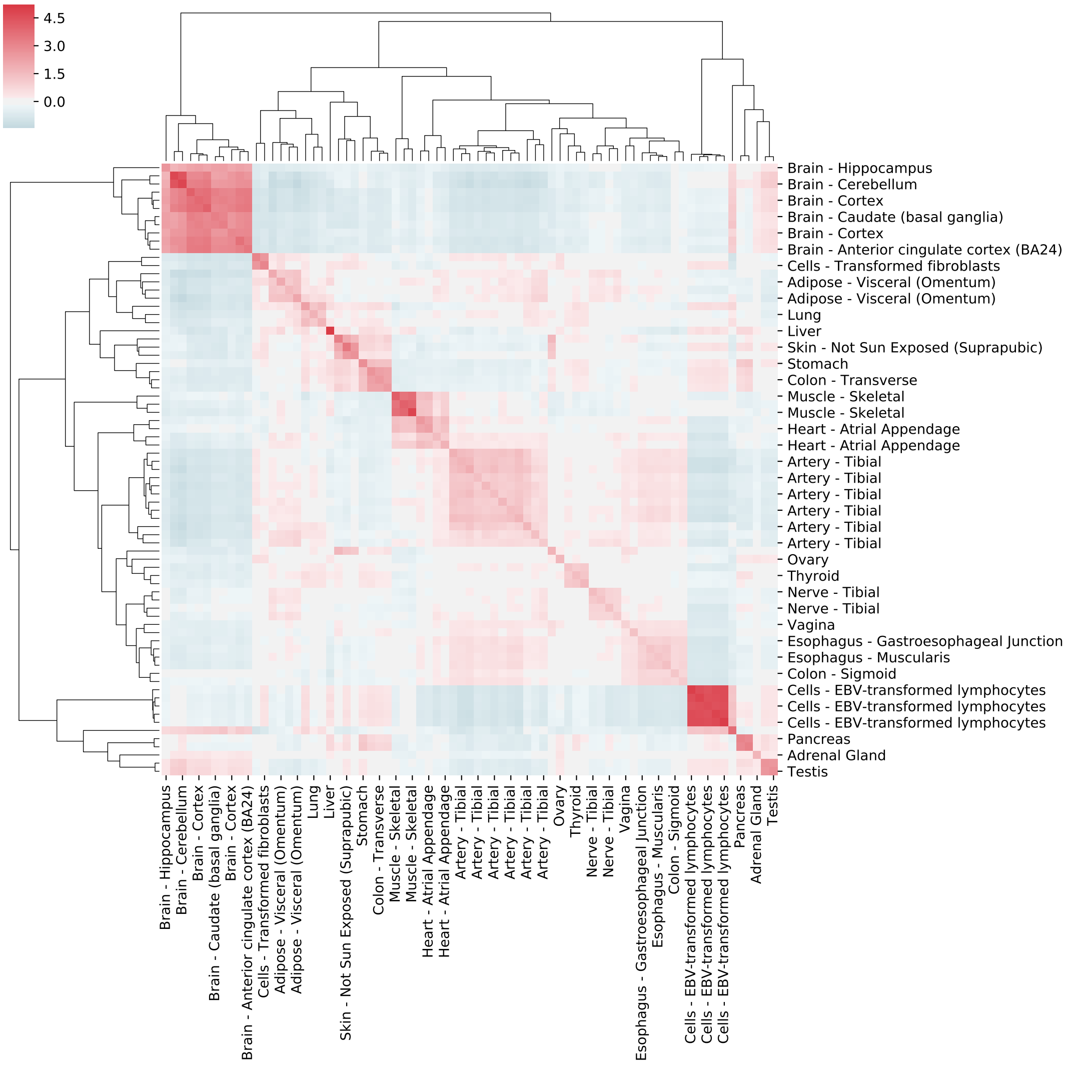


Figure 12: **H3K27ac: $\mathbf{\Sigma}_C$ for Bayesian Linear Mixed Model** Estimated noise $\mathbf{\Sigma}_C$ assuming dependence between the conditions for the GTEx dataset.

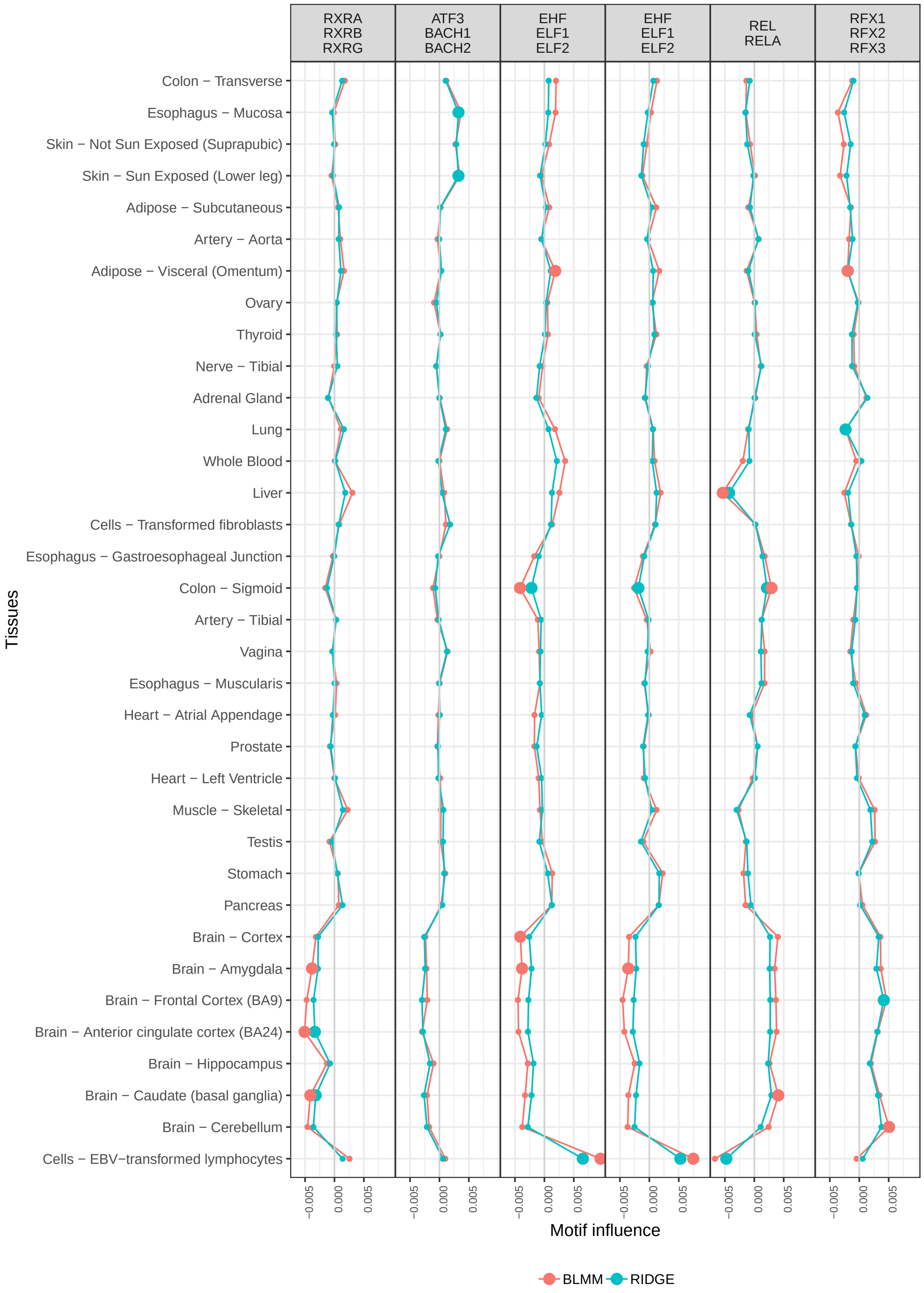


Figure 13: **GTEx: Most similar motif scores between methods** Motif values for the five highest correlated motif scores over all tissues.

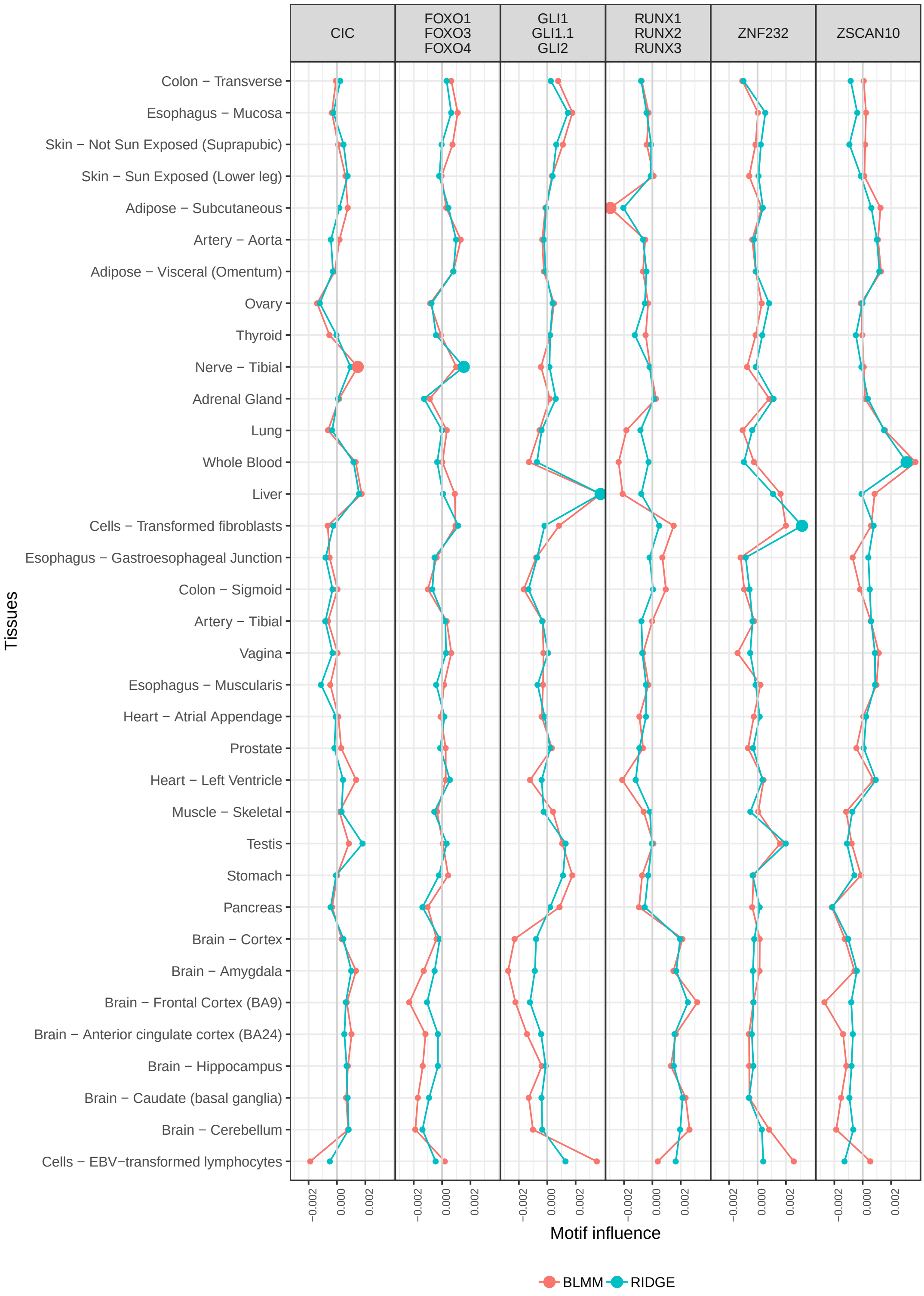


Figure 14: **GTEx: Least similar motif scores between methods** Motif values for the five lowest correlated motif scores over all tissues.

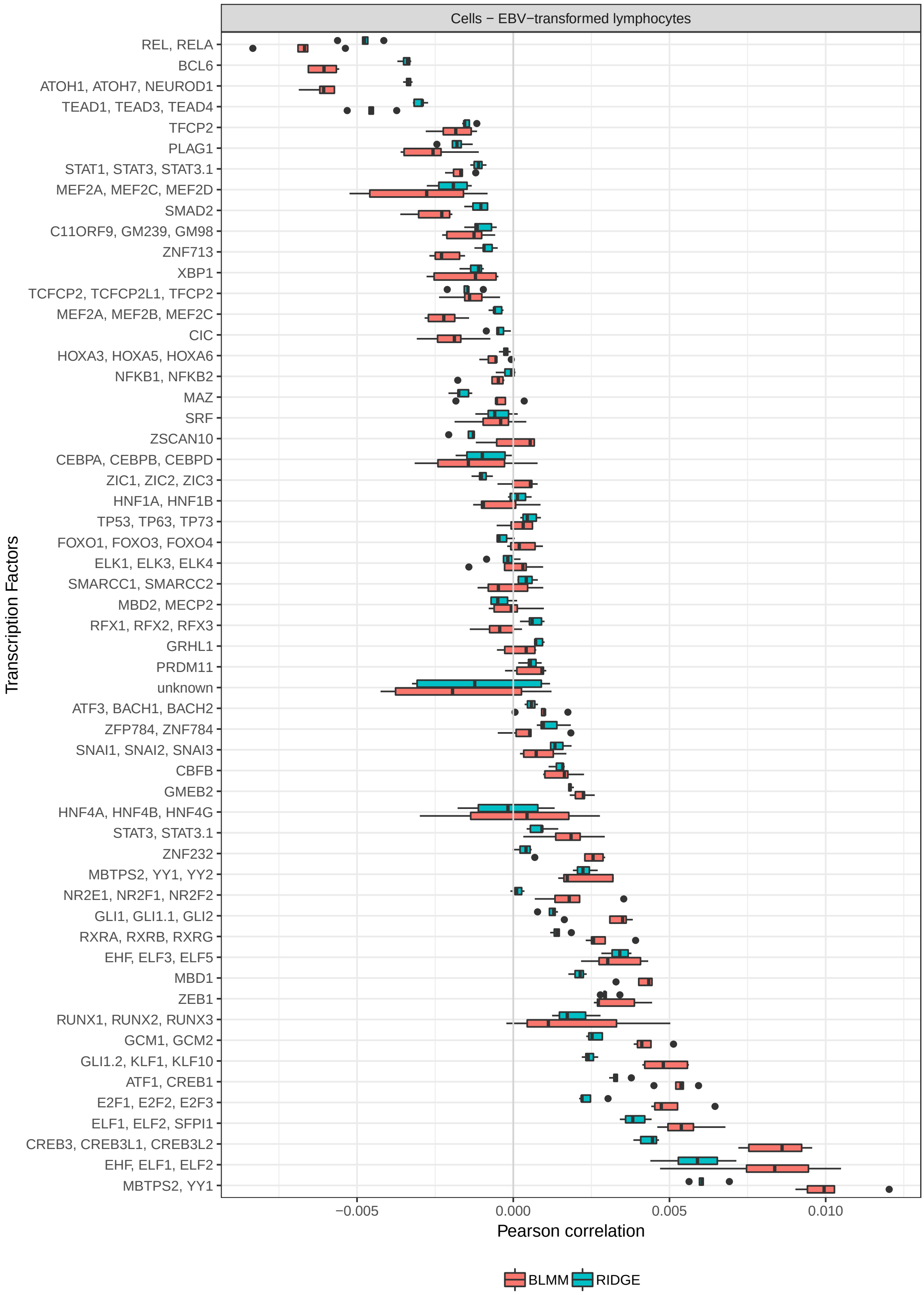


Figure 15: **GTEx: High variation between replicates** Variation of all chosen 56 motif scores on exemplary tissue EBV - cellline.
